# Riparian buffer management, rather than surrounding forest cover and buffer width, drives pest attacks in oil palm plantations

**DOI:** 10.64898/2026.02.10.704761

**Authors:** Li Yuen Chiew, Yehezkiel Jahuri, Sabidee Rizan, Arthur Y. C. Chung, Razy Japir, Tharaka S. Priyadarshana, Eleanor M. Slade

## Abstract

The rapid expansion of oil palm plantations in Southeast Asia has caused extensive deforestation and landscape fragmentation. Riparian buffers (vegetated strips along the edges of rivers) have been shown to enhance biodiversity, water quality, and erosion control. However, plantation managers have raised concerns that these buffers may harbour pests such as nettle caterpillars, bagworms, and rhinoceros beetles (*Oryctes rhinoceros*). These pests damage the palms and facilitate the spread *Ganoderma boninense* (a fungal pathogen). Using causal inference modelling we examined how riparian buffer characteristics (width and habitat quality), oil palm age, and surrounding landscape features influence pest and disease incidence in oil palms adjacent to riparian areas in Sabah, Malaysian Borneo. We surveyed 47,500 palms for pest and disease damage and used mark–release–recapture techniques to track *O. rhinoceros* movements in oil palms adjacent to riparian buffers. Most *O. rhinoceros* activity (66.30%) occurred within the plantations, and only 6.10% occurred within riparian buffers, with limited movement between habitats. Oil palm age was a dominant driver of pest attacks: young palms were more susceptible to lepidopteran caterpillars and *O. rhinoceros*, whereas *G. boninense* was more prevalent in mature palms. Neither the surrounding forest cover nor the quality of the riparian buffer affected the incidence of pest attacks. Riparian buffer width increased *O. rhinoceros* attacks, reduced *G. boninense* infection, and had no effect on lepidopteran caterpillars, highlighting that surrounding forest cover and riparian buffers do not drive pest attacks in oil palm plantations. Instead, management of oil palms within the buffers s is likely to be more important in managing pests; increases in invasive oil palms within the buffers increased the incidence of caterpillar damage, and higher numbers of remnant old oil palms increased *O. rhinoceros* attacks in adjacent oil palms. Overall, riparian buffers were found to contribute little to pest spillover, suggesting that their biodiversity and connectivity benefits outweigh minor pest risks, especially if invasive young and remnant old oil palms within the buffers are effectively managed and native vegetation restored.

## 1. INTRODUCTION

The conversion of natural habitats to agroecosystems in the tropics accounts for more than 70% of global deforestation and represents a major threat to biodiversity (Curtis et al., 2018). Within this broader pattern of agricultural expansion, oil palm (*Elaeis guineensis*) plantations have emerged as a dominant driver of forest loss (Meijaard et al., 2020). Although Southeast Asia, particularly Indonesia and Malaysia, dominates global oil palm production, cultivation is increasingly expanding into other tropical regions, raising concerns for biodiversity, ecosystem functioning, and human health (Ayompe et al., 2021; Meijaard et al., 2020). This expansion is driven by sustained global demand for palm oil, highlighting the challenge of balancing agricultural productivity with environmental sustainability.

In response, several certification schemes have been introduced to address the environmental and social impacts of palm oil production. These include the Roundtable on Sustainable Palm Oil scheme (RSPO; rspo.org/), and the Malaysian Sustainable Palm Oil (MSPO, www.mpocc.org.my/mspo-certification-scheme) and Indonesian Sustainable Palm Oil (ISPO, ispo-org.or.id) schemes, all of which aim to ensure that palm oil supplied to the global markets meets sustainability standards.

A key legal requirement in RSPO certification is the conservation of riparian buffers (i.e., vegetated strips along rivers), which play a vital role in stabilising riverbanks, reducing sedimentation, and limiting pollution entering watercourses (Lucey et al., 2018). Beyond these hydrological functions, riparian buffers provide a range of ecosystem services (Riis et al., 2020) and serve as important habitats for wildlife within oil palm landscapes (Bicknell et al., 2023; Deere et al., 2022). Wider buffers support higher species diversity (Deere et al., 2022), function as thermal refuges (Williamson et al., 2022), and facilitate animal movement across fragmented landscapes (Gray et al., 2019, 2022). However, many oil palm plantations in Southeast Asia were established prior to the introduction of sustainability guidelines, resulting in the absence of riparian buffers or the retention of narrow, highly degraded strips (Woodham et al., 2019). Even where buffers are present, their maintenance and ecological integrity are often compromised, and best-practice guidelines for buffer design and management have only recently been formalised (Lucey et al., 2018; Luke et al., 2020; Meijaard et al., 2018).

Despite their recognised ecological value, the oil palm industry has raised concerns that forested riparian buffers may facilitate the spread of pests and diseases into adjacent plantations (Luke et al., 2019). A key concern is that riparian buffers could serve as breeding habitats for rhinoceros beetles (*Oryctes rhinoceros* Linnaeus, 1758), a major pest of oil palm (Paudel et al., 2022). These beetles damage palms by feeding on fronds and attacking the apical meristem, which can lead to palm mortality (Egonyu et al., 2022). Moreover, it has been suggested that *O. rhinoceros* may contribute to the spread of *Ganoderma* fungus (*G. boninense*) by transporting fungal spores. Infection of the palm trunk with *G. boninense* causes basal stem rot and structural collapse of the palm (Hushiarian et al., 2013; Manjeri, 2014). Infestations of *O. rhinoceros* can reduce yields by up to 25% (Liau & Ahmad, 1991), while *G. boninense* infections may result in yield losses of 50–80% (Priwiratama & Susanto, 2014).

Several lepidopteran species also pose significant threats to oil palm production. Nettle caterpillars (Lepidoptera: Limacodidae) can reduce yields by up to 30% within two years of infestation (Potineni & Saravanan, 2013), while bagworms (Lepidoptera: Psychidae) may cause yield losses of up to 43% over a two-year period under high infestation levels (Kamarudin & Wahid, 2010). Despite these concerns, empirical evidence testing whether riparian buffers increase pest pressure in adjacent oil palm plantations remains limited.

This study addresses this knowledge gap by investigating (1) potential spillover of *O. rhinoceros* between riparian buffers and oil palm plantations, and (2) how riparian buffer characteristics (width and habitat quality), plantation attributes (palm age), and surrounding landscape features (forest cover and proximity to forests and riparian edges) influence pest and disease incidence. We combined a mark–release–recapture (MRR) experiment with causal inference approaches to test the following hypotheses (Figure 1):

**Figure 1.**
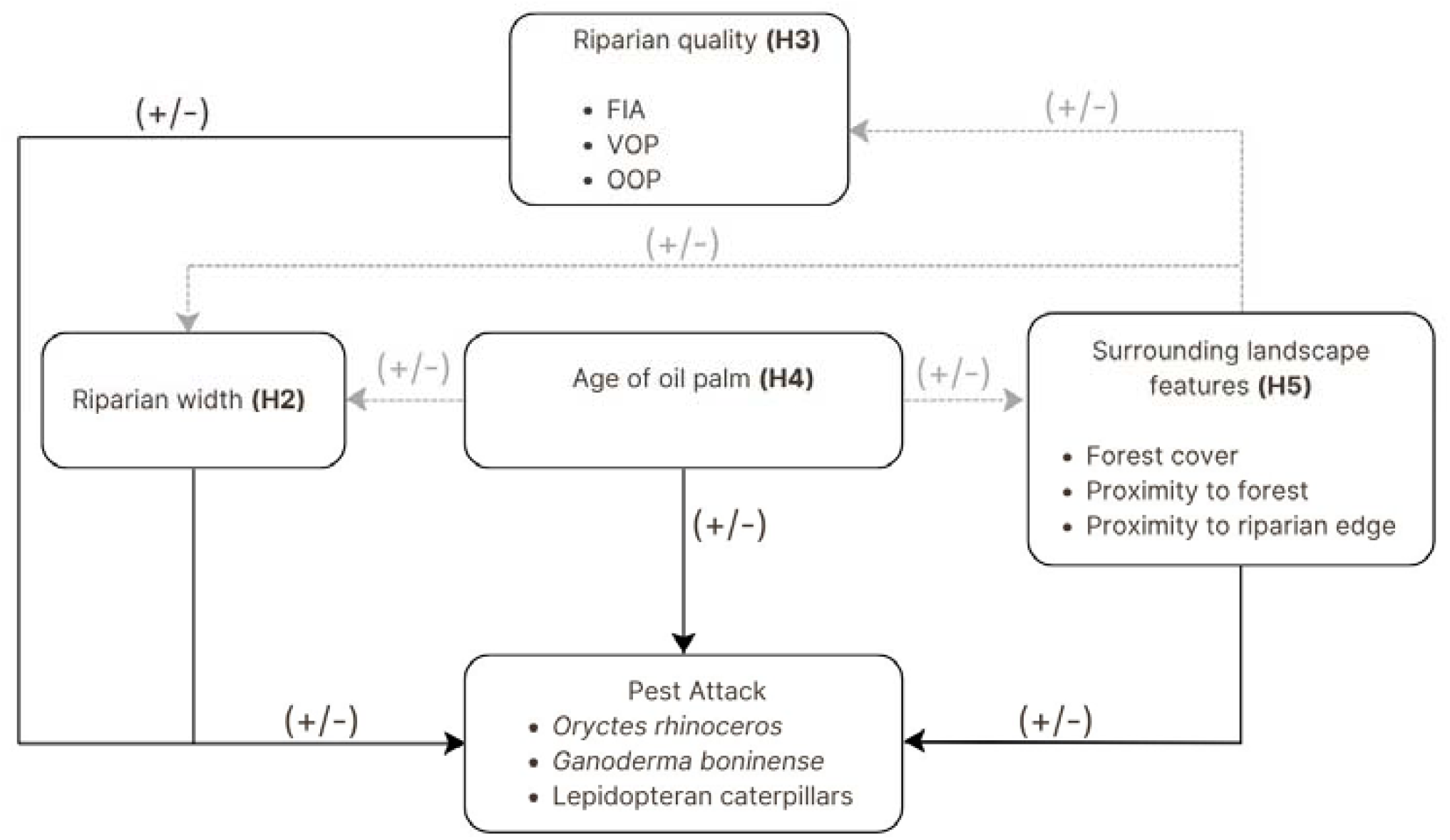
Graphical representation illustrating the hypothesized causal relationships in this study. Solid arrows represent hypothesised direct effects on pest attack, while dotted arrows indicate indirect effects mediated through intermediate variables (e.g. riparian buffer width or habitat quality). Plus (+) and minus (−) symbols denote the expected direction of effects, depending on ecological context. H2–H5 correspond to Hypotheses 2–5 tested in this study. See Introduction for detailed explanations.

### H1. Riparian buffers act as a source of Oryctes rhinoceros spillover into oil palm plantations

Following plantation managers’ concerns, we predicted that rhinoceros beetles move between riparian buffers and plantations, with greater net movement from buffers into the plantations, thereby increasing pest pressure.

### H2. Riparian buffer width has direct and indirect effects pest attacks in oil palm plantations

We hypothesised that wider buffers would provide resources such as deadwood and vegetation cover that can be exploited by pests (*O. rhinoceros*, lepidopteran caterpillars, and *G. boninense*), potentially increasing pest abundance or movement into plantations. Indirect effects may mediate through improved habitat quality or increased establishment of invasive oil palms within wider buffers (see *H3* below). Conversely, wider buffers may also support natural enemies that reduce pest impacts.

### H3. Riparian buffer habitat quality influences pest incidence in adjacent oil palm plantations

We hypothesised that higher-quality riparian buffers, characterised by greater forest integrity (measured by the Forest Integrity Assessment tool, see Methods below) and lower abundance of invasive and retained old oil palms, would either suppress pest attacks by supporting natural enemies or increase pest attacks if buffers provide suitable resources or breeding habitat for pests.

### H4. Oil palm age has direct and indirect effects on pest attacks

We predicted that younger palms are more susceptible to lepidopteran caterpillars and *O. rhinoceros* due to softer tissues and higher vulnerability (Beaudoin-Ollivier et al., 2017), whereas older palms experience different pest pressures, including higher susceptibility to fungal infections such as *G. boninense* (Jazuli et al., 2022). Palm age may also indirectly influence pest incidence through interactions with riparian buffer width and habitat quality, as plantation age affects vegetation structure, understory development, and microclimatic conditions.

### H5. Surrounding landscape features influence pest attacks directly and indirectly

Landscape attributes, including forest cover at multiple spatial scales (5–30 km radii), proximity to forested areas, and distance to riparian edge, may increase pest pressure if forests act as pest sources, as commonly assumed by the oil palm industry. Alternatively, forests may reduce pest attacks by supporting natural enemies. Landscape attributes may also indirectly influence pest incidence by shaping the ecological quality and integrity of riparian buffers.

## 2. MATERIALS AND METHODS

### 2.1. Study site

The study was conducted in RSPO-certified (rspo.org/) oil palm plantations (n =19 sites) in northeastern Sabah, Malaysian Borneo, between February and July 2022 (Figure 2). All plantations were owned and managed by Wilmar International Limited (www.wilmar-international.com/) and were located within three estates: Sapi I & II, Rumidi, and Reka Halus. The plantation area has a mean annual temperature of 26.50° C and receives approximately 2,497 mm of rainfall per year (Wilmar, internal records).

**Figure 2.**
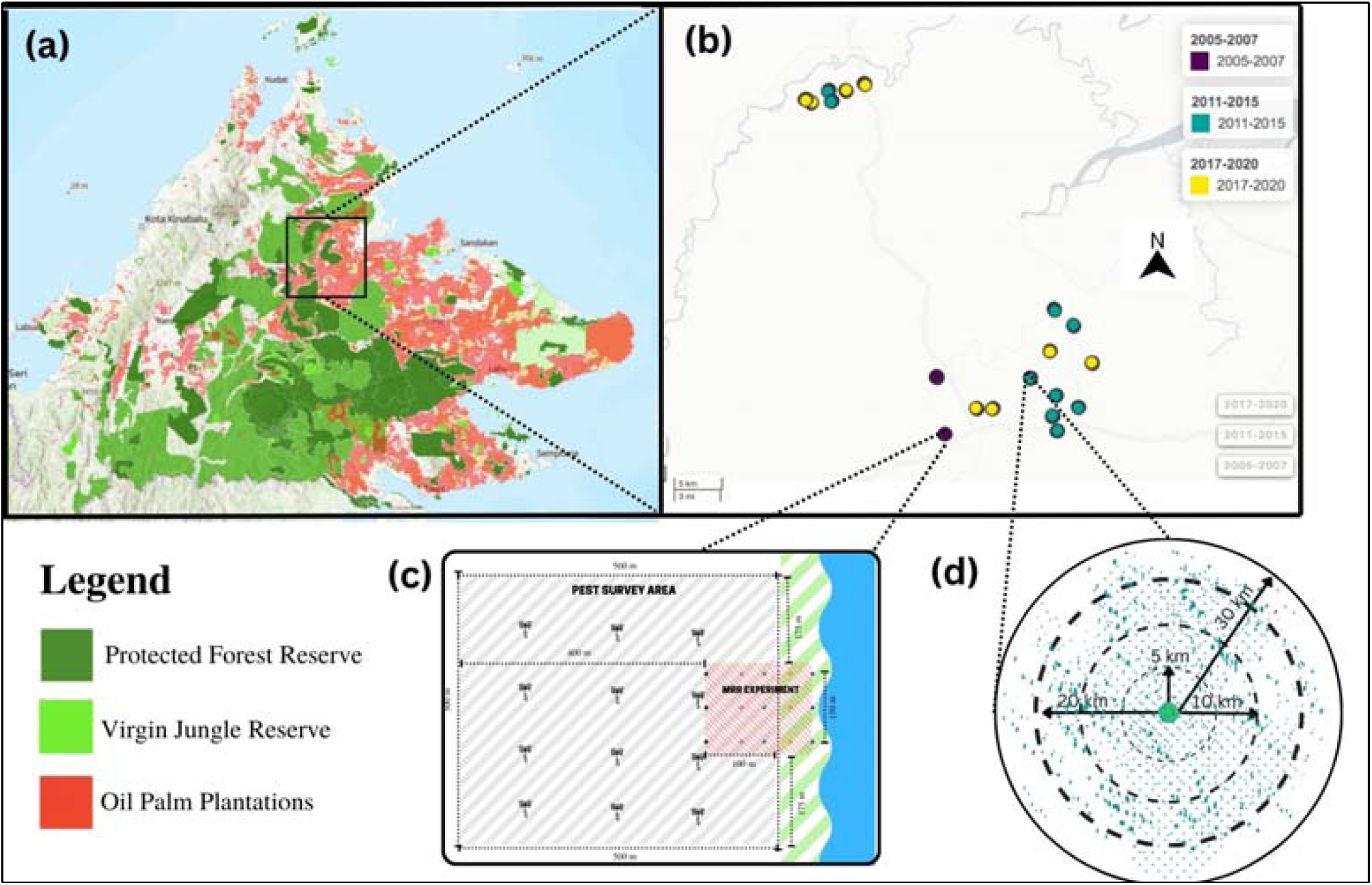
(a). Forest cover map of Sabah obtained from Hutanwatch.com (https://www.hutanwatch.com/sabah-land-use); (b) Locations of study sites in RSPO-certified oil palm plantations, Sabah, Malaysian Borneo (N=19 sites). The coloured circles show the locations of 19 study sites listed in Appendix Table S1, categorized by planting years; (c) Illustration of the (500m x 500m) pest survey area and Mark-Release-Recapture (MRR) experiment; (d) Percentage of forest cover in 5km, 10km, 20km, and 30km radius.

The plantations are located adjacent to the Bidu-bidu Forest Reserve, and most riparian areas within the estates are classified as “High Conservation Value (HCV)” zones (HCV 4), reflecting their importance for ecosystem services, such as watershed protection and erosion control (Wilmar International Conservation Initiatives, 2017; Brown & Senior, 2014). Each study site included a riparian buffer, with buffer widths ranging from 3 to 51 m (median = 16 m; Appendix: Table S1). Sites were separated by a minimum distance of 1.6 km (± 886 m) to ensure spatial independence (Figure 2b).

Oil palms across the sites represented a range of growth stages, including young palms (planted 2017–2020), mature fruiting palms (2011–2015), and old palms (2005–2007). Palms were planted at 10 m intervals, and management practices were standardised across all plantations. Ground cover was dominated by ferns and grass. Insecticides and herbicides were rarely used and were not applied during the study period.

### 2.2. Mark–release–recapture (MRR) experiment

Movement of *O. rhinoceros* beetles was assessed using pheromone traps baited with ethyl 4-methyloctnoate (E4-MO; SIME RB Pheromone), the standard monitoring compound for this pest (Paudel et al., 2023). The pheromone was suspended in a polymer sachet between a metal vane above a plastic bucket trap (Figure S1). To prevent overheating of the captured beetles, leaf material was placed inside traps located in open areas.

At each of the 19 sites, we deployed 12–18 traps (depending on riparian buffer width) arranged in a linear array extending from within the riparian buffer, across the buffer-plantation boundary, and into the oil palm plantation (Figure 3). This design follows established protocols for assessing invertebrate spillover in Sabah (Gray et al., 2016). Traps were positioned at the height of the palm fruiting zone (approximately 1.5 m above ground for mature palms), enabling capture of beetles feeding in the canopy as well as individuals emerging from decomposing empty fruit bunches and dead fronds used as mulch (Kamarudin & Wahid, 2004). The traps were operated for 24 hours per sampling event. Captured beetles were collected alive, sexed, and individually marked on the elytra using puncture markings (Unruh & Chauvin, 1993; Figure S1-S2). Beetles were then released at their capture points to track subsequent movement across the trap array. The MRR experiment continued until 10 marking-recapture events were completed. Capture points and dispersal paths were mapped using QGIS v3.28.1 (QGIS.org, 2022). For beetles that were recaptured multiple times each recapture was treated as an independent observation.

**Figure 3.**
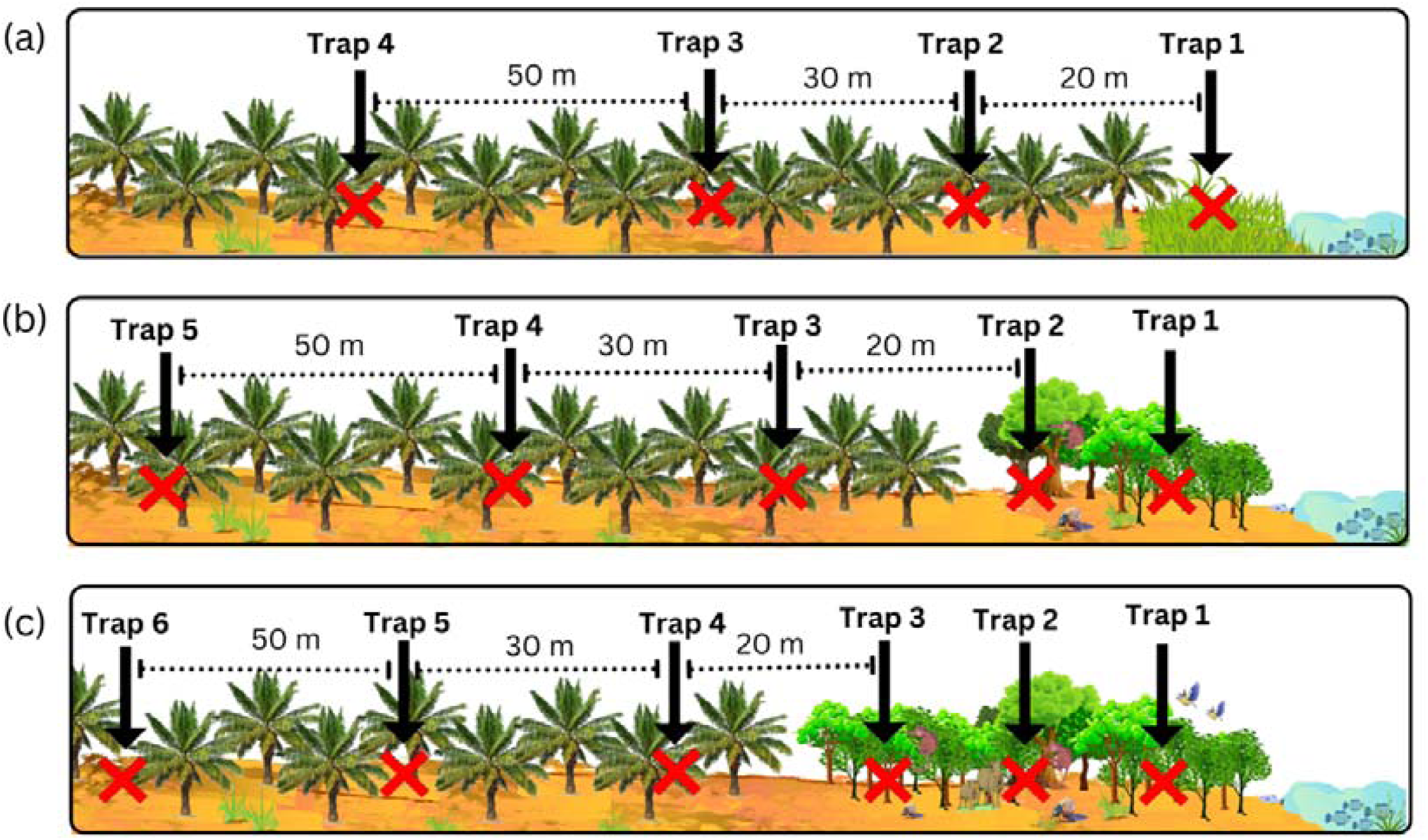
Schematic of the experimental design in three types of buffers as examples with the layout of sampling points for pheromone traps at **(a) oil palm with narrow riparian buffer** (< 15 metres); **(b) oil palm with minimum legal width** (15-25 metres); **(c) oil palm with wide riparian buffer** (> 25 metres). We established an array of 4-6 traps running along a transect from inside the riparian buffer, across the boundary, and into the plantation. There were three transects on each river, with 50 metres between each transect.

### 2.3. Visual surveys of pest attack

Pest damage was assessed in each site through visual surveys conducted by three trained observers (Figure 2c). Surveys targeted damage caused by lepidopteran caterpillars (nettle caterpillars *Setora sp.* and *Darna sp.*, and bagworms), *O. rhinoceros*, and *G. boninense*.

At each site, the survey covered a 500 × 500 m target area extending from the riparian buffer edge into the plantation (Figure 2c). Fifty parallel transects (500 m long) were established per site at 10 m intervals. Along each transect, 50 palms were assessed, resulting in 2,500 palms per site and 47,500 palms overall being assessed for pest attack. The same survey team conducted all assessments to ensure consistency; binoculars were used when required.

*O. rhinoceros* damage was identified by characteristic signs in the crown and fronds, including boring into petiole bases and central unopened leaves, fibrous frass within unopened fronds (Figure S3), and V-shaped cuts on fronds (Figure S4; Kenneth & John, 2020).

Lepidopteran caterpillar damage was identified based on defoliation patterns: mature caterpillars feed on the entire leaf blades, leaving only the midrib, whereas younger caterpillars scrape epidermal strips (Figures S5–S8). Because damage caused by nettle caterpillars and bagworms is difficult to distinguish visually, all lepidopteran damage was grouped together as “lepidopteran caterpillar attacks”.

*G. boninense* infections were identified by rotted roots, stem or trunk decay, and the presence of fungal fruiting bodies (Figure S9; Kamu et al., 2018). Additional indicators included wilting lower fronds that droop vertically to form a “skirt” (Figure S10; Chung, 2011).

### 2.4. Observed variables

To test our causal hypotheses (Figure 1), we measured eight variables describing riparian buffer characteristics, plantation attributes, and surrounding landscape context.

#### Riparian buffer variables

i. *Riparian buffer width –* measured as the distance (m) from the river’s edge to the plantation boundary. Tape measures were used for buffers < 20 m wide, while GPS coordinates were recorded for wider buffers and distances calculated as straight-line measurements.
ii. *Invasive young oil palms (VOPs) –* measured as the number of young oil palms established within the riparian buffer, counted along a 500 m transect within the buffer. These palms originate from dispersed fruit or from legacy plantings close to the river prior to RSPO certification.
iii. *Remnant old oil palms (OOPs) –* measured as the number of old, non-fruiting palms (poisoned or abandoned) within riparian buffers, counted along the same 500 m transects. These palms indicate buffer areas that were previously cleared and planted before buffer protection.
iv. *Riparian buffer quality –* assessed using the Forest Integrity Assessment (FIA) tool (Suggitt et al., 2021). Three assessors surveyed each site along a predefined transect (500 m long; 15–50 m wide depending on buffer width). The FIA consists of 50 questions covering forest structure, flora, fauna, hydrology, and disturbance. Scores range from 0 to 50, with higher values indicating higher forest integrity (Figure S11).

#### Plantation variable

v. *Oil palm age –* determined from plantation records and ranging from 4 to 19 years.

#### Surrounding landscape variables

vi. *Forest cover –* the percentage forest cover surrounding each site calculated at four spatial scales (5, 10, 20, and 30 km radii) using land-use maps from HUTANWATCH (www.hutanwatch.com/sabah-land-use).
vii. *Proximity to the forest –* the distance from each site to the nearest forest edge, measured using HUTANWATCH map tool.
viii. *Proximity to the riparian buffer* – the distance from the riparian edge to the location of each recorded pest attack recorded along the transects.

### 2.5. Data analysis

#### 2.5.1. Mark-release-recapture (MRR) analysis

We first used two-tailed binomial tests to assess whether marked *O. rhinoceros* recaptures showed a tendency to occur in oil palm plantations or riparian buffers, assuming an equal expected probability (50%) for each habitat (Weisburd et al., 2020). The empirical proportion represents the observed probability of recapture derived from the data, whereas the expected proportion represents the hypothesised probability under random movement. A non-significant result indicates that any deviation from the expected proportion could be attributed to random variation. In contrast, a significant result indicates that recaptures occurred at a frequency different from that expected under random movement, allowing inference on habitat preference or avoidance. To further evaluate the directionality of beetle movement (H1; see Introduction), we then applied one-tailed binomial tests to test whether movement between habitats was lower than expected, specifically from oil palm plantations to riparian buffers and from riparian buffers to oil palm plantations. These directional tests allowed us to determine whether beetle spillover in either direction occurred less frequently than predicted under random movement.

#### 2.5.2. Structural equation models

We first formulated multivariate causal hypotheses based on system-specific ecological knowledge (see *H2–H5* in the Introduction). These hypotheses encompassed both direct and indirect effects of riparian buffer characteristics, plantation attributes, and surrounding landscape context on pest attacks, resulting in interconnected variables in which some predictors act as both causes and effects. To formalise these hypothesised relationships, we constructed a directed acyclic graph (DAG) describing causal dependencies and conditional independencies among variables (see Figure 1; Grace et al., 2012). The DAG was then translated into a path model and evaluated using covariance-based Structural Equation Models (SEMs), which allow simultaneous estimation of multiple direct and indirect effects within a unified causal framework (Grace, 2006).

SEMs were fitted using the ‘sem’ function in the ‘lavaan’ package (Rosseel, 2012). Pest attack incidence was binary and therefore specified as an ordered categorical variable, with models estimated using the robust weighted least squares estimator (WLSMV), which is appropriate for categorical and non-normally distributed outcomes. To account for the hierarchical structure of the data, we adjusted statistical inference for clustering at the sampling-unit level using the ‘svydesign’ function in the ‘lavaan.survey’ package (Oberski, 2014). This procedure applies cluster-robust (sandwich) standard errors to the fitted SEM, ensuring that hypothesis tests and confidence intervals properly reflect the nested sampling design, while leaving the estimated path coefficients unchanged.

Model fit was assessed using the *χ²* statistic comparing observed and model-implied covariance matrices (Shipley, 2016). Path coefficients were standardised to facilitate comparison among effect strengths across predictors. Indirect effects were calculated as the product of coefficients along constituent paths, and total effects were obtained by summing direct and indirect effects (Appendix: Tables S2–S13).

Separate SEMs were fitted for lepidopteran caterpillars, *O. rhinoceros*, and *G. boninense*. For each pest, models were evaluated at four spatial scales corresponding to forest cover within 5, 10, 20, and 30 km radii. Model support across spatial scales was compared using AICc. The 10 km scale provided the best fit (lowest AICc) for all pests; results for this scale are therefore presented in the main text (Table 2).

**Table 1.** Observed variables and their variation (± standard deviation).

| Observed variables | Mean ± SD | Minimum | Maximum |
| --- | --- | --- | --- |
| Age of palms (years) | 7.21 ± 4.24 | 3 | 17 |
| Forest cover (percentage) |  |  |  |
| • 5 km scale | 1.25 ± 1.95 | 0 | 4.8 |
| • 10 km scale | 0.75 ± 1.14 | 0 | 2.7 |
| • 20 km scale | 3.12 ± 1.55 | 1.2 | 6.1 |
| • 30 km scale | 4.82 ± 1.39 | 2.2 | 7.3 |
| Proximity to the forest (km) | 3.35 ± 2.28 | 0.65 | 8.05 |
| Proximity to the riparian buffer (m) | 250 ± 145 | 0 | 495 |
| Riparian width (m) | 17.53 ± 11.22 | 3 | 51 |
| Riparian quality (FIA) (scores) | 24.5 ± 5.07 | 15 | 34 |
| Invasive young oil palms (VOPs) (no.) | 28.74 ± 38.17 | 0 | 150 |
| Remnant old oil palms (OOPs) (no.) | 10 ± 24.76 | 0 | 83 |

**Table 2.** The best model explaining pest attack within different scales of forest cover percentage. SEMs – RB = The model fitted with the *O. rhinoceros* attack. SEMs – GF = The model fitted with the *G. boninense* attack. SEMs – CP = The model fitted with the lepidopteran caterpillar attack. AICs = Akaike information criterion values for each model corrected for small sample sizes. The best scale that has the lowest AICc value is shown in bold.

| Model | Forest Cover Scale | AICc |
| --- | --- | --- |
| SEMs - RB | 5 km | -106118.46 |
|  | **10 km** | **-150468.8** |
|  | 20 km | -93750.08 |
|  | 30 km | -136974.49 |
| SEMs - GF | 5 km | -330893 |
|  | **10 km** | **-375249.8** |
|  | 20 km | -318535.6 |
|  | 30 km | -361741.7 |
| SEMs - CP | 5 km | -67203.33 |
|  | **10 km** | **-111544.63** |
|  | 20 km | -54812.22 |
|  | 30 km | -98101.43 |

To aid ecological interpretation, selected SEM effects on pest attack were expressed as absolute changes in predicted probability (Bollen, 1989). Predicted probabilities were derived from the model-implied link function using the estimated threshold for the binary outcome and the linear predictor. Because predictor variables were standardized, a baseline linear predictor value of zero corresponds to mean conditions across covariates. For each predictor of interest, predicted probabilities were calculated at this baseline and following a one standard deviation increase in the focal variable, holding all other variables constant. The difference between these probabilities was reported as a percentage-point change in the probability of pest attack, providing an interpretable measure of effect magnitude for practitioners. Statistical significance was assessed at alpha 0.05. All analyses were conducted in R v4.4.1 (www.r-project.org).

## 3. RESULTS

### 3.1. Mark–release–recapture (MRR) experiment

#### H1. Riparian buffers act as a source of Oryctes rhinoceros beetle spillover into oil palm plantations

Across the 19 study sites, a total of 1,261 *O. rhinoceros* beetles were marked, with 211 recapture events recorded, yielding an overall recapture rate of 16.70% (Table S1). Of these recapture events, 166 involved beetles recaptured more than once. Among all marked beetles, 77.30% were initially trapped in oil palm plantations and 22.70% in riparian buffers. Of the marked recaptured beetles, 66.30% were recaptured in oil palm plantations, whereas only 6.10% were recaptured within riparian buffers.

Movement between habitats occurred in both directions: 14.70% of beetles moved from riparian buffers into oil palm plantations, while 12.90% moved from plantations into riparian buffers (Figure 4). Most recaptures occurred between 2 and 20 days after release, with only 0.70% of individuals recaptured within 24 hours. On average, beetles moved 108.70 m over an eight-day period, although one individual travelled 1.60 km between sites within 24 hours.

**Figure 4.**
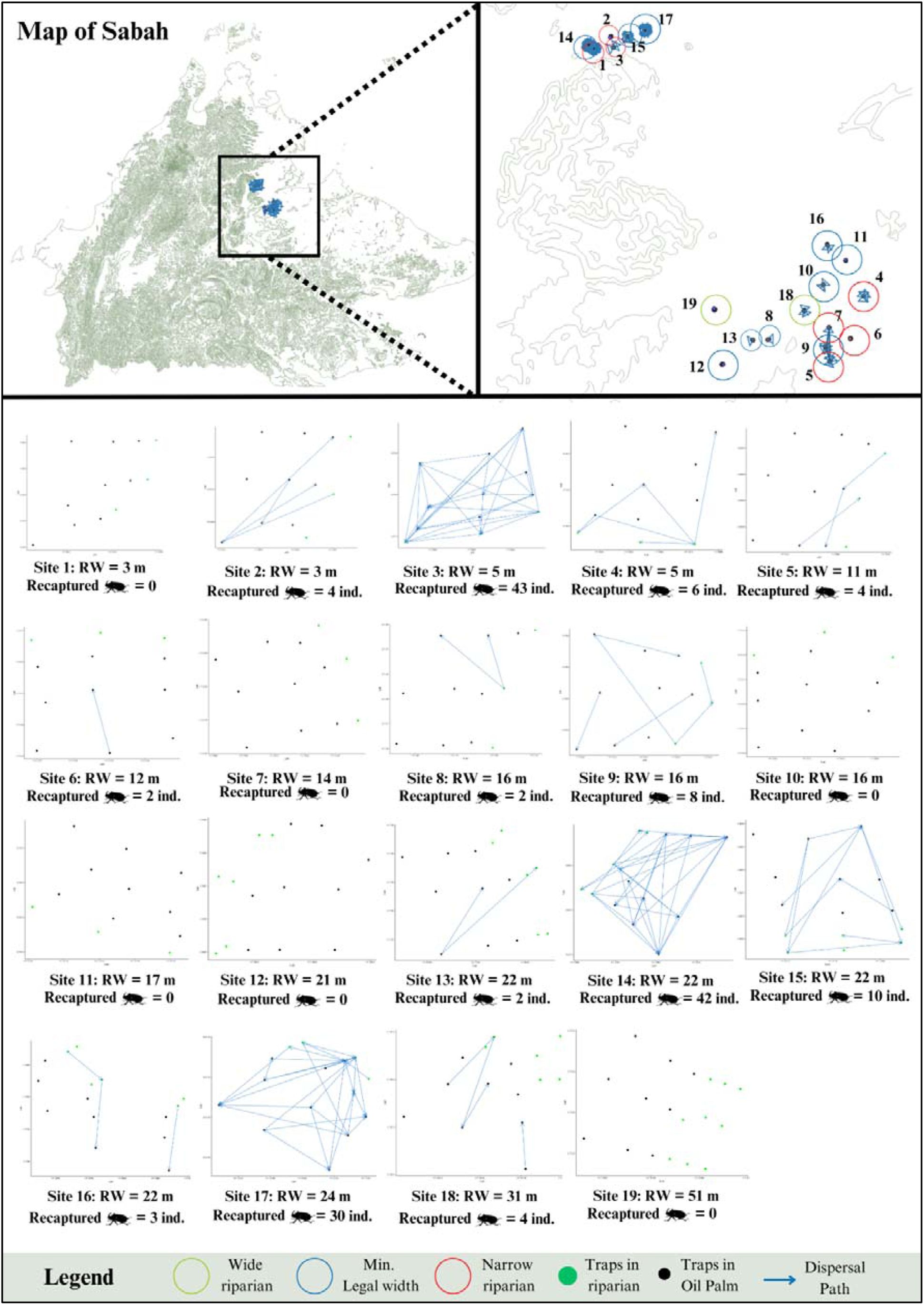
Annotated diagram of the sites showing overall and close-up views of the Mark-Release-Recapture (MRR) movement paths.

A significantly higher proportion of marked beetles were recaptured in oil palm plantation traps than in riparian buffer traps (N = 211, expected proportion [ExPr] = 0.50, empirical proportion [EmPr] = 0.77, 95% confident intervals [CIs] = 0.71, 1.00, *P* < 0.001). Beetles marked in oil palm plantations showed no tendency was to move into riparian buffers (N = 211, ExPr = 0.50, EmPr = 0.10, 95% CIs = 0.060, 1.00, *P* < 0.001). Similarly, beetles marked within riparian buffers were unlikely to move into plantations (N = 211, ExPr = 0.5, EmPr = 0.11, 95% CIs = [0.074, 1.00], *P* < 0.001).

### 3.2. Effects of riparian buffers, plantation attributes, and surrounding landscape features on pest attacks

#### H2. Riparian buffer width has direct and indirect positive effects on pest attacks in oil palm plantations

We detected pest-specific direct effects of riparian buffer width, but no consistent indirect effects, on pest attack incidence (Figure 5; Table S2-S13). Riparian buffer width had a direct positive effect on *O. rhinoceros* attack incidence (Figure 5a; Table 3; Supp Table S3): a one standard-deviation (SD) increase in buffer width was associated with a 1.28 percentage-point increase in the predicted probability of *O. rhinoceros* attack. In contrast, buffer width had a direct negative effect on *G. boninense* infection (Figure 5b; Table 3; Supp Table S7), with the predicted probability of infection decreasing by 0.011 percentage-points per one SD increase in buffer width. Riparian buffer width had no direct effect on lepidopteran caterpillar attack incidence (Figure 5c; Table 3; Supp Table S11). Although buffer width positively influenced riparian habitat quality (FIA scores), this did not translate into strong indirect effects of buffer width on any of the studied pest groups via habitat quality or other mediating variables (Figure 5; Supp Table S3, S7, S11).

**Figure 5.**
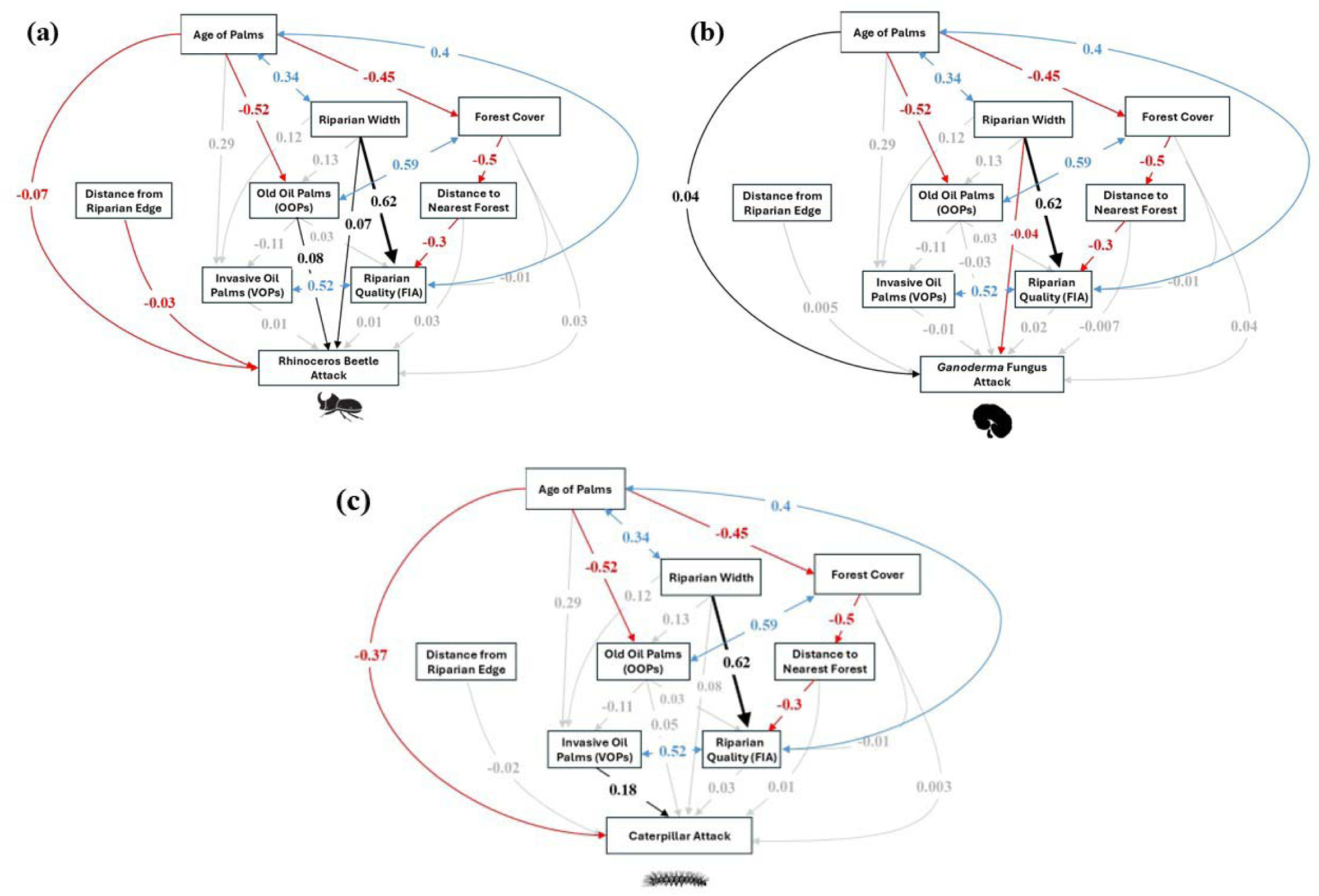
Path diagrams illustrating direct and indirect causal links between observed variables at the 10km forest cover scale for (a) *O. rhinoceros* attack; (b) *G. boninense* attack; (c) lepidopteran caterpillar attack. Standardized direct effect sizes are indicated along the arrows, representing the direction of causal links and proportional to the standardized effect sizes. Black arrows indicate significantly positive effects, red arrows indicate significantly negative effects, blue arrows indicate correlated effects, and lightly shaded arrows indicate non-significant effects at alpha level of 0.05. All panels share identical global fit statistics (χ^2^ = 6.86; df =12; p-values: 0.867) because the models are based on the same causal structure and conditional independence (d-separation) set and were fitted to the same dataset; pest-specific differences are reflected in path coefficients and indirect effects rather than overall model fit. For detailed statistics, including indirect and total effects between observed variables, see Appendix Table S5-S16.

**Table 3.** Predicted percentage-point change in the probability of pest group attack associated with a one standard deviation (SD) increase in each predictor. Estimates were derived from a probit structural equation model fitted in lavaan with cluster-robust standard errors (lavaan.survey). Probability changes were calculated using the estimated outcome threshold (τ) from the underlying lavaan model and standardized regression coefficients. Positive values indicate increased probability of infection, whereas negative values indicate reduced probability. Standard errors (SE), z-values, and p-values are based on the cluster-corrected model.

| Predictor variable | Standardized coefficients | Standard Errors (SE) | Percentage-point change* | P-value |
| --- | --- | --- | --- | --- |
| *O. rhinoceros attack* |  |  |  |  |
| • Riparian width | 0.065 | 0.035 | 1.28 | 0.061 |
| • FIA | -0.013 | 0.022 | -0.24 | 0.573 |
| • VOP | 0.011 | 0.026 | 0.21 | 0.664 |
| • OOP | 0.084 | 0.025 | 1.66 | **0.001** |
| • Proximity to the forest | -0.025 | 0.031 | -0.46 | 0.421 |
| • Proximity to the riparian buffer | -0.026 | 0.012 | -0.48 | **0.026** |
| • Forest cover | 0.032 | 0.035 | 0.62 | 0.355 |
| • Oil palm age | -0.072 | 0.030 | -1.30 | **0.016** |
| *G. boninense attack* |  |  |  |  |
| • Riparian width | -0.040 | 0.014 | -0.01 | **0.005** |
| • FIA | 0.017 | 0.017 | 0.01 | 0.303 |
| • VOP | -0.012 | 0.013 | -0.00 | 0.353 |
| • OOP | -0.026 | 0.017 | -0.01 | 0.117 |
| • Proximity to the forest | -0.007 | 0.011 | -0.00 | 0.519 |
| • Proximity to the riparian buffer | 0.005 | 0.003 | 0.00 | 0.078 |
| • Forest cover | 0.042 | 0.018 | 0.01 | **0.018** |
| • Oil palm age | 0.038 | 0.013 | 0.01 | **0.004** |
| *Lepidopteran caterpillar attack* |  |  |  |  |
| • Riparian width | 0.075 | 0.047 | 2.96 | 0.115 |
| • FIA | 0.031 | 0.105 | 1.21 | 0.771 |
| • VOP | 0.176 | 0.070 | 7.01 | **0.011** |
| • OOP | 0.051 | 0.066 | 2.01 | 0.442 |
| • Proximity to the forest | 0.013 | 0.066 | 0.51 | 0.845 |
| • Proximity to the riparian buffer | -0.023 | 0.015 | -0.90 | 0.122 |
| • Forest cover | 0.003 | 0.075 | 0.14 | 0.963 |
| • Oil palm age | -0.367 | 0.093 | -13.78 | **<0.001** |

\* Percentage-point (pp) change in pest attack probability providing an interpretable measure of effect magnitude for practitioners. It refers to how much the probability shifts on a 0–100 scale (e.g., from 44% to 47% is +3 pp).

#### H3. Riparian buffer habitat quality influences pest incidence in adjacent oil palm plantations

Contrary to our hypothesis, riparian habitat quality (FIA scores) showed no direct or indirect effect on lepidopteran caterpillar attacks, *O. rhinoceros* attacks, or *G. boninense* infection (Figure 5; Supp Table S3, S7, S11).

In contrast, the abundance of invasive young oil palms (VOPs) within riparian buffers had a direct positive effect on lepidopteran caterpillar attack incidence (Figure 5c; Table 3; Supp Table S11): a one SD increase in VOP abundance was associated with a 7.01 percentage-point increase in the predicted probability of lepidopteran caterpillar attack. VOP abundance had no direct effects on *O. rhinoceros* or *G. boninense* attack incidence. However, VOPs exerted indirect effects on *O. rhinoceros* attack incidence, primarily mediated through FIA, with a one SD increase in VOP abundance associated with a 0.21 percentage-point increase in the predicted probability of *O. rhinoceros* attack (Table 3; Supp Table S3).

The abundance of remnant old oil palms (OOPs) had a direct positive effect on of *O. rhinoceros* attack incidence (Figure 5a; Table 3; Supp Table S3): a one SD increase in OOP abundance increased the predicted probability of attack by 1.66 percentage points. No direct or indirect effects of OOP abundance were detected for lepidopteran caterpillar attacks or *G. boninense* infection (Supp Table S7).

#### H4. Oil palm age has direct and indirect effects on pest attacks

Oil palm age had direct negative effects on *O. rhinoceros* and lepidopteran caterpillar attack incidence (Figure 5a, c; Table 3; Supp Table S3, S11). A one SD increase in palm age reduced the predicted probability of lepidopteran caterpillar attack by 13.78 percentage points and O. rhinoceros attack by 1.30 percentage points. In contrast, palm age had a direct positive effect on *G. boninense* infection (Figure 5b; Table 3; Supp Table S7), with the predicted probability of infection increasing by 0.012 percentage points per one SD increase in palm age. No indirect effects of palm age on any of the studied pest groups were detected.

#### H5. Surrounding landscape features influence pest attacks directly and indirectly

Contrary to our hypothesis, neither surrounding forest cover nor distance to the nearest forest had direct or indirect effects on lepidopteran caterpillar attacks, *O. rhinoceros* attacks, or *G. boninense* infection (Figure 5; Supp Table S5–S16). However, there is a direct effect of proximity to riparian edge to *O. rhinoceros* attacks, with the predicted probability of infection decreasing by 0.48 percentage points per one SD increase in the distance from riparian edge (Table 3; Supp Table S3). There was an also a direct but insignificant effect of proximity to riparian edge on lepidopteran caterpillar attack with a one SD increase in the distance associated with a 0.90 percentage-point decrease in the predicted probability of lepidopteran caterpillar attack (Table 3; Supp Table S11). No direct or indirect effect in proximity to riparian edge was detected in *G. boninense* infection (Table 3; Supp Table S7).

## 4. DISCUSSION

Conserving natural habitat patches within and around agricultural landscapes is widely recognised as essential for maintaining biodiversity and ecosystem services (Cole et al., 2020; Deere et al., 2022; Graziano et al., 2022). In oil palm systems, riparian buffers are a key conservation intervention intended to protect waterways, maintain habitat connectivity, and support ecological functioning (Deere et al., 2022; Luke et al., 2020). However, concerns persist within the oil palm industry that vegetated riparian buffers may exacerbate pest pressure in adjacent plantations. By explicitly testing these assumptions across multiple pest groups (i.e., *Oryctes rhinoceros*, *Ganoderma boninense,* and lepidopteran caterpillars) using a combination of mark–release–recapture experiments and causal inference models, our study provides empirical evidence that riparian buffers alone do not increase pest attack incidence in certified oil palm plantations.

We found that *O. rhinoceros* beetles predominantly moved within plantations rather than between riparian buffers and plantations, and that pest incidence was more strongly associated with palm age and the presence of young invasive oil palms within buffers than with buffer width or surrounding forest cover. While O. *rhinoceros* attack probability increased slightly near wider buffers, wider buffers were associated with reduced *G. boninense* infection, and no relationship was detected between buffer width and lepidopteran caterpillar attacks. Overall, our results suggest that pest dynamics are driven more by plantation structure and buffer condition than by the presence and amount of riparian vegetation.

### 4.1. Riparian buffers are not a major source of *Oryctes rhinoceros* spillover

Contrary to common assumptions among plantation managers, our mark-release-recapture experiment found little evidence that riparian buffers act as a major source of *O. rhinoceros* spillover into plantations. Although beetles were occasionally captured within riparian buffers, most captured and recaptured individuals occurred within plantations, and cross-habitat movements were infrequent. Average movement distances were short (108.7 m on average within 24 h), and recaptures typically occurred several days after release (between 2-20 days), indicating limited dispersal between habitats.

These movement patterns suggest that *O. rhinoceros* primarily inhabits and exploits plantation habitats rather than forested riparian areas, and there is little and active dispersal between habitats. This behaviour is consistent with the ecology of *O. rhinoceros*, which relies heavily on decomposing organic material for oviposition and larval development (Oyewale & Makinde, 2023). Within plantations resources such as dead trunks, fallen fronds, and empty fruit bunches provide abundant and continuous breeding substrates (Jackson et al., 2020), whereas such materials are typically less abundant in riparian and forest vegetation.

These findings suggest that pest management strategies targeting *O. rhinoceros* should primarily focus on reducing breeding resources within plantations themselves, rather than viewing riparian buffers and adjacent forest patches as primary pest sources.

### 4.2. Riparian buffer width and quality have pest-specific effects

Our hypotheses predicted that wider riparian buffers could either increase pest incidence by providing additional resources (e.g., decaying wood and organic debris) or reduce pest pressure by supporting natural enemies. The results support neither outcome universally. Instead, the effects of riparian buffers were pest-specific and mediated by buffer structure and condition, and particularly the presence of invasive young, or remnant old, oil palms within the buffer.

Riparian buffer width showed small but contrasting effects across pest groups, increasing *O. rhinoceros* attack probability while reducing *G. boninense* infection, and having no effect on lepidopteran caterpillars. These contrasting responses highlight that buffer effects cannot be generalised across pest taxa. For insect pests such as *O. rhinoceros*, wider buffers may increase the likelihood of attacks on palms adjacent to riparian edges, potentially due to edge effects or the accumulation of organic residues near buffer boundaries (Jackson et al., 2020). In contrast, wider buffers were associated with lower *G. boninense* incidence, providing no evidence that riparian vegetation acts as a source of *Ganoderma* outbreaks in adjacent oil palm.

Importantly, riparian habitat quality, as measured by the Forest Integrity Assessment (FIA), did not directly influence pest incidence. In contrast, the presence of invasive young oil palms (VOPs) and remnant old oil palms (OOPs) within buffers showed clear associations with pest attacks. VOPs were strongly linked to increased lepidopteran caterpillar attack incidence, likely because they act as host plants that sustain herbivore populations within the riparian buffer. OOPs were associated with increased *O. rhinoceros* attacks, potentially because decaying palm material from these provides breeding sites for beetles within the buffer. Together, these findings indicate that degraded riparian buffers dominated by invasive young or remnant old oil palms may increase pest habitat suitability, whereas buffers restored with native vegetation may minimise pest risks while maintaining ecological benefits.

### 4.3. Younger oil palm plantations are more vulnerable to insect pests

Palm age emerged as one of the strongest predictors of pest incidence in our study. Younger oil palm plantations (< 4 years) were significantly more susceptible to *O. rhinoceros* and lepidopteran caterpillar attacks than older plantations. This pattern is consistent with previous research showing that young palms are more vulnerable due to their tender foliage, rapid growth, and limited structural defences (Egonyu et al., 2022; Kenneth & John, 2020; Manjeri, 2014). Herbivorous pests are therefore most damaging during the early establishment phase, when palms are least resilient to defoliation and meristem damage.

In contrast, *G. boninense* infection was more prevalent in mature palms. This likely reflects the slow and progressive nature of fungal colonisation, with visible symptoms such as fruiting bodies appearing only at later stages of infection (Jazuli et al., 2022). Thus, early-stage infections in younger palms may have gone undetected, potentially underestimating true pathogen prevalence. Previous studies have also shown that plantations undergoing second or subsequent replanting cycles face elevated risks of *Ganoderma* outbreaks due to pathogen persistence in soil and residual root material (Snaddon et al., 2013). These contrasting age-related patterns highlight the need for age-specific pest and disease management strategies within oil palm plantations.

### 4.4. Limited influence of surrounding landscape features on pest attack incidence

Contrary to industry concerns, broader landscape features such as forest cover and proximity to forest did not influence pest incidence in our study. This suggests that surrounding forests are unlikely to act as major sources of pests for adjacent oil palm plantations. Similar findings have been reported in oil palm and other tropical agricultural systems, where pest pressure is often more strongly driven by local plantation management and habitat conditions than by landscape-scale forest cover (Denan et al., 2020; Gómez Mateus et al., 2023). Instead, our results indicate that pest dynamics are primarily shaped by within-plantation processes and conditions at the plantation-riparian interface.

We observed higher incidences of *O. rhinoceros* and lepidopteran caterpillar attacks near riparian edges. This pattern is likely linked to management practices at plantation margins rather than landscape context per se. In oil palm systems, organic residues such as empty fruit bunches, felled trunks, and decomposing palm material are often stockpiled near plantation edges for logistical convenience. This materials provide favourable breeding substrates for beetle larvae, which are known to proliferate in decaying organic matter (Manjeri, 2014). This suggests that elevated pest incidence near riparian edges reflects management-induced resource heterogeneity, not pest spillover from riparian buffers.

## 5.0 Management implications and future research directions

This study provides important insights into how pest dynamics in oil palm landscapes are shaped by interacting plantation-level and riparian management factors, rather than by the mere presence of adjacent forested habitats. In particular, our findings demonstrate that pest incidence is more strongly influenced by oil palm age and management of invasive or remnant oil palms within riparian buffers than by surrounding forest cover or the conservation of forest areas *per se* within plantations. These results directly address long-standing industry concerns that biodiversity conservation measures, such as riparian buffers, inevitably increase pest pressure in adjacent oil palm stands.

Although current sustainability guidelines and certification schemes recommend maintaining 20–100 m of natural vegetation along rivers (Lucey et al., 2018), our results indicate that buffer quality is at least as important as buffer width in shaping pest outcomes. Riparian buffers dominated by invasive young oil palms (VOPs) or remnant old oil palms (OOPs) were associated with higher pest incidence, particularly in younger plantations that are most vulnerable to damage. In contrast, riparian buffers with fewer invasive young or remnant old oil palms and greater native vegetation cover are likely to reduce pest habitat suitability while continuing to deliver key ecosystem services such as erosion control, water regulation, and habitat connectivity.

Based on these findings, we recommend that riparian management strategies move beyond a sole focus on buffer width and incorporate active restoration and maintenance of buffer quality. This includes the systematic removal of invasive young and retained old oil palms from riparian zones, avoidance of planting oil palms close to riparian edges, and replanting of native tree and understory species. Integrating pest management objectives into riparian buffer design and restoration guidelines would allow plantations to better balance agricultural productivity with biodiversity conservation goals, rather than treating these objectives as competing priorities.

Our results also show minimal spillover of *O. rhinoceros* beetles from riparian buffers into oil palm plantations, indicating that riparian conservation does not inherently conflict with maintaining high-yield production. This finding challenges the perception that forested buffers act as major pest reservoirs and supports the compatibility of riparian protection with effective pest control. Where beetle attacks increase near riparian edges, practical management interventions, such as maintaining a narrow vegetation-free gap between riparian buffers and oil palm rows or limiting the accumulation of organic residues near buffer boundaries, may help reduce local pest pressure without compromising buffer function. Future research should build on these findings by explicitly testing the mechanistic pathways linking riparian vegetation composition, microclimate, and pest-natural enemy interactions. Long-term monitoring across plantation age gradients, coupled with experimental riparian restoration trials, would help determine how improvements in buffer quality influences pest suppression and natural enemy abundance over time. Such studies are essential for refining evidence-based guidelines that integrate riparian conservation into sustainable pest management frameworks, ultimately enhancing both ecological resilience and agricultural sustainability in oil palm-dominated landscapes.

## ACKNOWLEDGMENTS

This project was funded by the Ministry of Education, Singapore, through the Academic Research Fund Tier 1 (Project No. RG119/19) to Associate Prof. Eleanor M. Slade. The research was conducted under the Sabah Biodiversity Centre (SaBC) access licence entitled “Riparian Protection versus Pest and Disease Control in Oil Palm Plantations” (JKM/MBS.1000-2/2 JLD.13 [119]). We thank Wilmar International Limited for granting access to their plantations (Sapi/Terusan/SekarImej/Sabahmas/SEARRP/001), as well as for providing accommodation and logistical support during fieldwork. Field data collection was supported by the South East Asia Rainforest Research Programme (SEARRP). We also thank Ong Xin Rui for helpful discussions on the data analysis.

---

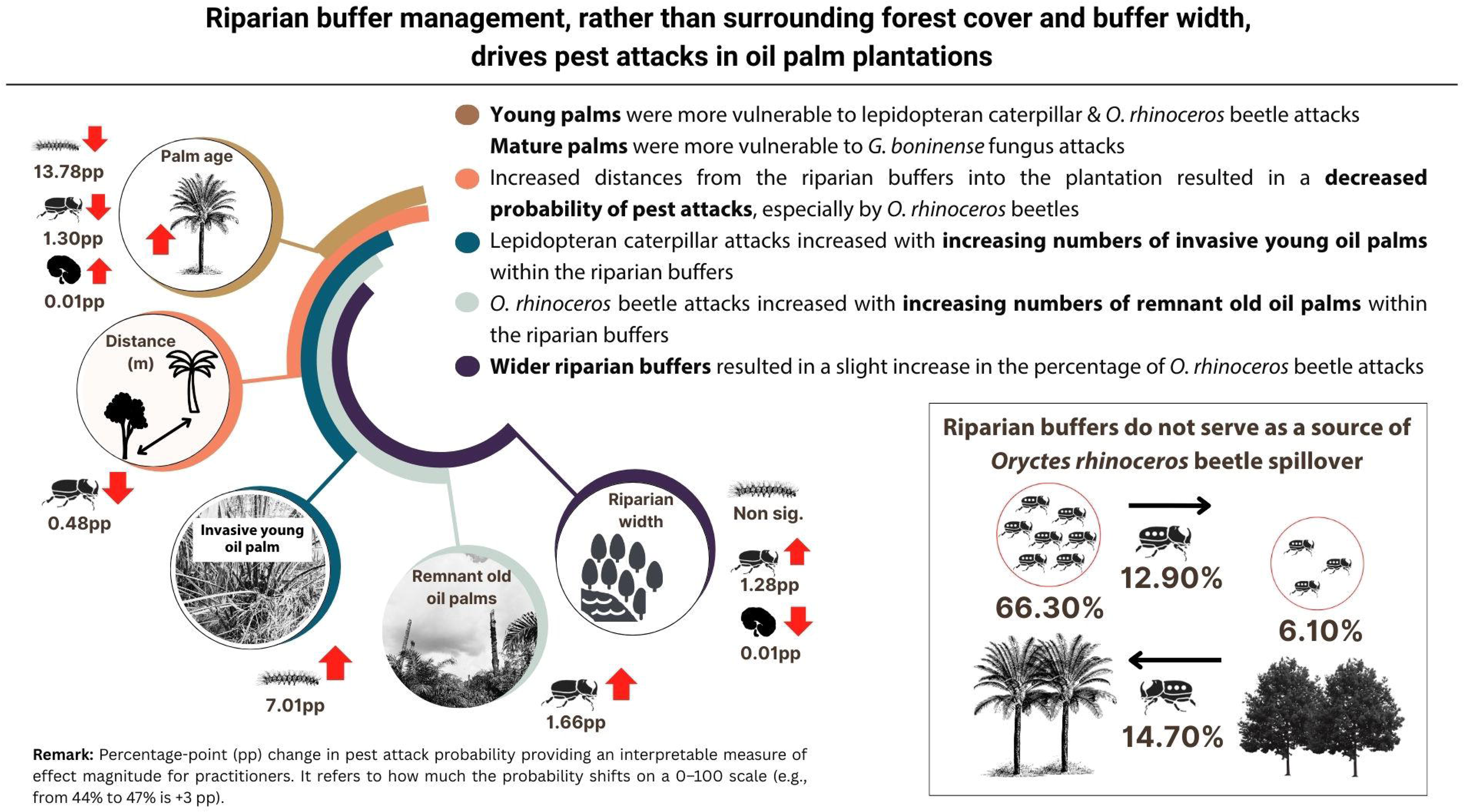

