## Supplementary materials for "Riparian buffer management, rather than surrounding forest cover and buffer width, drives pest attacks in oil palm plantations"

**Supplementary Information**

**Table S1** Summary of Mark-Release-Recapture (MRR) data.

| **Sites** | **Planting**  **Year** | **Riparian width** | **Within Riparian** | | | **Within OP** | | | **Total** | |
| --- | --- | --- | --- | --- | --- | --- | --- | --- | --- | --- |
|  |  | **(m)** | **Marked** | **Recaptured** | **Recaptured (%)** | **Marked** | **Recaptured** | **Recaptured (%)** | **Marked** | **Recaptured** |
| 1 | 2013 | 3 | 0 | 0 | 0.00 | 30 | 0 | 0.00 | 30 | 0 |
| 2 | 2013 | 3 | 0 | 0 | 0.00 | 27 | 5 | 18.52 | 27 | 5 |
| 3 | 2020 | 5 | 0 | 0 | 0.00 | 277 | 57 | 20.58 | 277 | 57 |
| 4 | 2018 | 5 | 0 | 0 | 0.00 | 57 | 6 | 10.53 | 57 | 6 |
| 5 | 2013 | 11 | 17 | 1 | 5.88 | 32 | 4 | 12.50 | 49 | 5 |
| 6 | 2015 | 12 | 3 | 1 | 33.33 | 16 | 1 | 6.25 | 19 | 2 |
| 7 | 2014 | 14 | 0 | 0 | 0.00 | 22 | 0 | 0.00 | 22 | 0 |
| 8 | 2017 | 16 | 11 | 0 | 0.00 | 13 | 2 | 15.38 | 24 | 2 |
| 9 | 2011 | 16 | 26 | 2 | 7.69 | 18 | 6 | 33.33 | 44 | 8 |
| 10 | 2018 | 16 | 11 | 2 | 18.18 | 21 | 5 | 23.81 | 32 | 7 |
| 11 | 2014 | 17 | 8 | 0 | 0.00 | 15 | 0 | 0.00 | 23 | 0 |
| 12 | 2005 | 21 | 9 | 0 | 0.00 | 7 | 0 | 0.00 | 16 | 0 |
| 13 | 2019 | 22 | 9 | 1 | 11.11 | 16 | 1 | 6.25 | 25 | 2 |
| 14 | 2020 | 22 | 46 | 11 | 23.91 | 282 | 40 | 14.18 | 328 | 51 |
| 15 | 2019 | 22 | 28 | 6 | 21.43 | 77 | 8 | 10.39 | 105 | 14 |
| 16 | 2014 | 22 | 27 | 2 | 7.41 | 10 | 2 | 20.00 | 37 | 4 |
| 17 | 2019 | 24 | 30 | 20 | 66.67 | 68 | 20 | 29.41 | 98 | 40 |
| 18 | 2012 | 31 | 4 | 1 | 25.00 | 27 | 3 | 11.11 | 31 | 4 |
| 19 | 2007 | 51 | 6 | 0 | 0.00 | 8 | 0 | 0.00 | 14 | 0 |

**Table S2** Direct and indirect effects between observed variables in the proposed causal model at **5 km scale forest cover *O. rhinoceros* attack**. For more details, see Figure A3. Statistically significant (alpha level of 0.05) effects/coefficients are in bold letters. b = Unstandardized effects. b* = Standardised effects / coefficients. SE = Standard error. p = p-value.

|  |  | **Direct effect** | | | | **Indirect effect** | | | | **Total effect** | | | |
| --- | --- | --- | --- | --- | --- | --- | --- | --- | --- | --- | --- | --- | --- |
| **Affected variable** | **Causal variable** | ***b*** | ***SE*** | ***b**** | ***p*** | ***b*** | ***SE*** | ***b**** | ***p*** | ***b*** | ***SE*** | ***b**** | ***p*** |
| Rhinoceros beetle attack | Riparian width | 0.018 | 0.007 | 0.062 | 0.014 | -0.015 | 0.016 | -0.045 | 0.333 | 0.003 | 0.012 | 0.017 | 0.825 |
| Rhinoceros beetle attack | Riparian quality | -0.089 | 0.139 | -0.014 | 0.524 | 0.006 | 0.004 | -0.000 | 0.149 | -0.083 | 0.139 | -0.014 | 0.548 |
| Rhinoceros beetle attack | VOPs | 0.010 | 0.020 | 0.012 | 0.613 | -0.048 | 0.022 | -0.056 | 0.034 | -0.037 | 0.020 | -0.044 | 0.068 |
| Rhinoceros beetle attack | OOPs | 0.104 | 0.030 | 0.080 | 0.001 | 0.019 | 0.015 | 0.067 | 0.194 | 0.123 | 0.025 | 0.147 | 0.000 |
| Rhinoceros beetle attack | Distance to nearest forest | -0.038 | 0.043 | -0.027 | 0.373 | -0.037 | 0.042 | -0.058 | 0.378 | -0.075 | 0.028 | -0.085 | 0.008 |
| Rhinoceros beetle attack | Distance from riparian edge | -0.056 | 0.026 | -0.026 | 0.032 | 0.000 | 0.000 | 0.000 | 0.613 | -0.056 | 0.026 | -0.026 | 0.032 |
| Rhinoceros beetle attack | Forest cover | 0.056 | 0.060 | 0.034 | 0.351 | 0.038 | 0.029 | 0.107 | 0.183 | 0.094 | 0.046 | 0.141 | 0.040 |
| Rhinoceros beetle attack | Age of palms | -0.053 | 0.021 | -0.069 | 0.012 | -0.042 | 0.023 | -0.044 | 0.066 | -0.095 | 0.027 | -0.113 | 0.000 |
| Riparian quality | Forest cover | 0.004 | 0.062 | 0.017 | 0.945 | 0.034 | 0.024 | -0.160 | 0.150 | 0.038 | 0.060 | -0.143 | 0.525 |
| Riparian quality | Distance to nearest forest | -0.065 | 0.029 | -0.295 | 0.028 | -0.001 | 0.033 | 0.137 | 0.972 | -0.066 | 0.045 | -0.158 | 0.142 |
| VOPs | OOPs | -0.171 | 0.170 | -0.111 | 0.314 | -0.076 | 0.050 | -0.149 | 0.124 | -0.247 | 0.162 | -0.260 | 0.127 |
| Riparian quality | OOPs | 0.003 | 0.038 | 0.013 | 0.946 | 0.001 | 0.006 | -0.126 | 0.863 | 0.004 | 0.033 | -0.113 | 0.915 |
| OOPs | Riparian width | 0.016 | 0.038 | 0.07 | 0.680 | -0.045 | 0.048 | -0.170 | 0.351 | -0.029 | 0.056 | -0.100 | 0.603 |
| VOPs | Riparian width | 0.041 | 0.064 | 0.123 | 0.512 | 0.045 | 0.045 | 0.108 | 0.327 | 0.086 | 0.043 | 0.231 | 0.047 |
| Riparian quality | Riparian width | 0.028 | 0.005 | 0.624 | 0.000 | -0.002 | 0.002 | -0.028 | 0.471 | 0.026 | 0.005 | 0.596 | 0.000 |
| OOPs | Age of palms | -0.292 | 0.153 | -0.501 | 0.055 | 0.002 | 0.006 | 0.024 | 0.708 | -0.290 | 0.152 | -0.477 | 0.056 |
| VOPs | Age of palms | 0.259 | 0.146 | 0.288 | 0.076 | 0.056 | 0.064 | 0.094 | 0.379 | 0.315 | 0.119 | 0.382 | 0.008 |

**Table S3** Direct and indirect effects between observed variables in the proposed causal model at **10 km scale forest cover *O. rhinoceros* attack**. For more details, see Figure A3. Statistically significant (alpha level of 0.05) effects/coefficients are in bold letters. b = Unstandardized effects. b* = Standardised effects / coefficients. SE = Standard error. p = p-value.

|  |  | **Direct effect** | | | | **Indirect effect** | | | | **Total effect** | | | |
| --- | --- | --- | --- | --- | --- | --- | --- | --- | --- | --- | --- | --- | --- |
| **Affected variable** | **Causal variable** | ***b*** | ***SE*** | ***b**** | ***p*** | ***b*** | ***SE*** | ***b**** | ***p*** | ***b*** | ***SE*** | ***b**** | ***p*** |
| Rhinoceros beetle attack | Riparian width | 0.019 | 0.007 | 0.065 | 0.010 | -0.014 | 0.017 | -0.040 | 0.403 | 0.005 | 0.012 | 0.025 | 0.700 |
| Rhinoceros beetle attack | Riparian quality | -0.081 | 0.140 | -0.013 | 0.562 | 0.009 | 0.012 | 0.001 | 0.459 | -0.073 | 0.139 | -0.012 | 0.602 |
| Rhinoceros beetle attack | VOPs | 0.010 | 0.020 | 0.011 | 0.633 | -0.048 | 0.023 | -0.054 | 0.032 | -0.039 | 0.021 | -0.043 | 0.059 |
| Rhinoceros beetle attack | OOPs | 0.108 | 0.029 | 0.084 | 0.000 | 0.019 | 0.015 | 0.065 | 0.188 | 0.128 | 0.024 | 0.149 | 0.000 |
| Rhinoceros beetle attack | Distance to nearest forest | -0.035 | 0.045 | -0.025 | 0.433 | -0.098 | 0.112 | -0.057 | 0.383 | -0.133 | 0.078 | -0.082 | 0.091 |
| Rhinoceros beetle attack | Distance from riparian edge | -0.056 | 0.026 | -0.026 | 0.032 | 0.000 | 0.000 | 0.000 | 0.633 | -0.056 | 0.026 | -0.026 | 0.032 |
| Rhinoceros beetle attack | Forest cover | 0.091 | 0.098 | 0.032 | 0.350 | 0.041 | 0.045 | 0.101 | 0.358 | 0.133 | 0.069 | 0.133 | 0.054 |
| Rhinoceros beetle attack | Age of palms | -0.055 | 0.020 | -0.072 | 0.007 | -0.042 | 0.022 | -0.040 | 0.056 | -0.097 | 0.027 | -0.112 | 0.000 |
| Riparian quality | Forest cover | -0.006 | 0.109 | -0.013 | 0.959 | 0.065 | 0.041 | -0.103 | 0.113 | 0.060 | 0.106 | -0.116 | 0.572 |
| Riparian quality | Distance to nearest forest | -0.066 | 0.028 | -0.304 | -0.019 | 0.007 | 0.108 | 0.133 | 0.951 | -0.060 | 0.109 | -0.171 | 0.585 |
| VOPs | OOPs | -0.171 | 0.171 | -0.112 | 0.318 | -0.079 | 0.050 | -0.142 | 0.118 | -0.250 | 0.163 | -0.254 | 0.125 |
| Riparian quality | OOPs | 0.005 | 0.037 | 0.027 | 0.883 | 0.001 | 0.005 | -0.124 | 0.913 | 0.006 | 0.032 | -0.097 | 0.852 |
| OOPs | Riparian width | 0.030 | 0.039 | 0.130 | 0.459 | -0.047 | 0.051 | -0.175 | 0.358 | -0.017 | 0.061 | -0.045 | 0.773 |
| VOPs | Riparian width | 0.041 | 0.064 | 0.122 | 0.517 | 0.043 | 0.047 | 0.103 | 0.364 | 0.084 | 0.044 | 0.225 | 0.053 |
| Riparian quality | Riparian width | 0.027 | 0.005 | 0.616 | 0.000 | -0.001 | 0.002 | -0.022 | 0.568 | 0.026 | 0.005 | 0.594 | 0.000 |
| OOPs | Age of palms | -0.304 | 0.154 | -0.517 | 0.048 | 0.004 | 0.007 | 0.044 | 0.547 | -0.300 | 0.153 | -0.473 | 0.050 |
| VOPs | Age of palms | 0.259 | 0.148 | 0.287 | 0.080 | 0.058 | 0.067 | 0.095 | 0.389 | 0.317 | 0.118 | 0.382 | 0.007 |

**Table S4** Direct and indirect effects between observed variables in the proposed causal model at **20 km scale forest cover *O. rhinoceros* attack**. For more details, see Figure A3. Statistically significant (alpha level of 0.05) effects/coefficients are in bold letters. b = Unstandardized effects. b* = Standardised effects/ coefficients. SE = Standard error. p = p-value.

|  |  | **Direct effect** | | | | **Indirect effect** | | | | **Total effect** | | | |
| --- | --- | --- | --- | --- | --- | --- | --- | --- | --- | --- | --- | --- | --- |
| **Affected variable** | **Causal variable** | ***b*** | ***SE*** | ***b**** | ***p*** | ***b*** | ***SE*** | ***b**** | ***p*** | ***b*** | ***SE*** | ***b**** | ***p*** |
| Rhinoceros beetle attack | Riparian width | 0.014 | **0.007** | 0.050 | 0.049 | -0.012 | 0.014 | -0.033 | 0.397 | 0.003 | 0.011 | 0.017 | 0.807 |
| Rhinoceros beetle attack | Riparian quality | 0.012 | 0.174 | 0.002 | 0.945 | -0.002 | 0.007 | -0.027 | 0.731 | 0.010 | 0.170 | -0.025 | 0.955 |
| Rhinoceros beetle attack | VOPs | 0.001 | 0.019 | 0.001 | 0.949 | -0.048 | 0.024 | -0.040 | 0.045 | -0.047 | 0.024 | -0.038 | 0.057 |
| Rhinoceros beetle attack | OOPs | 0.131 | 0.033 | 0.101 | 0.000 | 0.014 | 0.011 | 0.023 | 0.191 | 0.145 | 0.029 | 0.124 | 0.000 |
| Rhinoceros beetle attack | Distance to nearest forest | -0.059 | 0.026 | -0.042 | 0.022 | 0.036 | 0.034 | 0.011 | 0.293 | -0.023 | 0.049 | -0.031 | 0.638 |
| Rhinoceros beetle attack | Distance from riparian edge | -0.056 | 0.026 | -0.026 | 0.032 | 0.000 | 0.000 | 0.000 | 0.949 | -0.056 | 0.026 | -0.026 | 0.032 |
| Rhinoceros beetle attack | Forest cover | 0.067 | 0.050 | 0.032 | 0.178 | -0.022 | 0.032 | -0.012 | 0.479 | 0.045 | 0.050 | 0.020 | 0.371 |
| Rhinoceros beetle attack | Age of palms | -0.050 | 0.022 | -0.066 | 0.024 | -0.039 | 0.021 | -0.034 | 0.064 | -0.089 | 0.025 | -0.100 | 0.000 |
| Riparian quality | Forest cover | -0.066 | 0.055 | -0.193 | 0.235 | -0.026 | 0.026 | -0.219 | 0.312 | -0.092 | 0.054 | -0.412 | 0.090 |
| Riparian quality | Distance to nearest forest | -0.051 | 0.030 | -0.223 | 0.086 | -0.033 | 0.039 | -0.113 | 0.400 | -0.084 | 0.040 | -0.336 | 0.037 |
| VOPs | OOPs | -0.171 | 0.170 | -0.111 | 0.313 | -0.076 | 0.050 | -0.150 | 0.128 | -0.247 | 0.161 | -0.261 | 0.126 |
| Riparian quality | OOPs | -0.002 | 0.020 | -0.006 | 0.949 | -0.004 | 0.003 | -0.195 | 0.113 | -0.006 | 0.020 | -0.201 | 0.778 |
| OOPs | Riparian width | 0.013 | 0.038 | 0.061 | 0.727 | -0.044 | 0.048 | -0.169 | 0.355 | -0.031 | 0.053 | -0.108 | 0.557 |
| VOPs | Riparian width | 0.042 | 0.063 | 0.123 | 0.511 | 0.045 | 0.044 | 0.109 | 0.311 | 0.087 | 0.044 | 0.232 | 0.049 |
| Riparian quality | Riparian width | 0.027 | 0.005 | 0.573 | 0.000 | 0.001 | 0.002 | 0.026 | 0.475 | 0.028 | 0.005 | 0.599 | 0.000 |
| OOPs | Age of palms | -0.291 | 0.151 | -0.497 | 0.054 | 0.002 | 0.007 | 0.020 | 0.755 | -0.289 | 0.150 | -0.477 | 0.055 |
| VOPs | Age of palms | 0.259 | 0.146 | 0.288 | 0.076 | 0.056 | 0.062 | 0.094 | 0.372 | 0.315 | 0.119 | 0.382 | 0.008 |

**Table S5** Direct and indirect effects between observed variables in the proposed causal model at **30 km scale forest cover *O. rhinoceros* attack**. For more details, see Figure A3. Statistically significant (alpha level of 0.05) effects/coefficients are in bold letters. b = Unstandardized effects. b* = Standardised effects/ coefficients. SE = Standard error. p = p-value.

|  |  | **Direct effect** | | | | **Indirect effect** | | | | **Total effect** | | | |
| --- | --- | --- | --- | --- | --- | --- | --- | --- | --- | --- | --- | --- | --- |
| **Affected variable** | **Causal variable** | ***b*** | ***SE*** | ***b**** | ***p*** | ***b*** | ***SE*** | ***b**** | ***p*** | ***b*** | ***SE*** | ***b**** | ***p*** |
| Rhinoceros beetle attack | Riparian width | 0.017 | 0.008 | 0.060 | 0.035 | -0.017 | 0.014 | -0.051 | 0.246 | 0.001 | 0.010 | 0.010 | 0.956 |
| Rhinoceros beetle attack | Riparian quality | -0.144 | 0.162 | -0.023 | 0.375 | 0.012 | 0.012 | 0.011 | 0.353 | -0.132 | 0.155 | -0.013 | 0.395 |
| Rhinoceros beetle attack | VOPs | 0.018 | 0.021 | 0.021 | 0.394 | -0.049 | 0.025 | -0.062 | 0.048 | -0.031 | 0.022 | -0.041 | 0.154 |
| Rhinoceros beetle attack | OOPs | 0.120 | 0.041 | 0.093 | 0.004 | 0.014 | 0.013 | 0.043 | 0.285 | 0.134 | 0.035 | 0.136 | 0.000 |
| Rhinoceros beetle attack | Distance to nearest forest | -0.026 | 0.035 | -0.019 | 0.451 | -0.052 | 0.077 | -0.015 | 0.495 | -0.079 | 0.078 | -0.034 | 0.315 |
| Rhinoceros beetle attack | Distance from riparian edge | -0.056 | 0.026 | -0.026 | 0.032 | 0.000 | 0.000 | 0.000 | 0.394 | -0.056 | 0.026 | -0.026 | 0.032 |
| Rhinoceros beetle attack | Forest cover | -0.075 | 0.072 | -0.032 | 0.301 | -0.001 | 0.033 | -0.005 | 0.985 | -0.075 | 0.072 | -0.037 | 0.297 |
| Rhinoceros beetle attack | Age of palms | -0.070 | 0.030 | -0.092 | 0.021 | -0.017 | 0.029 | -0.013 | 0.561 | -0.086 | 0.027 | -0.105 | 0.002 |
| Riparian quality | Forest cover | -0.108 | 0.059 | -0.29 | 0.069 | -0.028 | 0.046 | -0.207 | 0.542 | -0.136 | 0.044 | -0.497 | 0.002 |
| Riparian quality | Distance to nearest forest | -0.024 | 0.040 | -0.105 | 0.548 | -0.122 | 0.078 | -0.290 | 0.116 | -0.146 | 0.063 | -0.395 | 0.020 |
| VOPs | OOPs | -0.171 | 0.169 | -0.111 | 0.312 | -0.073 | 0.051 | -0.159 | 0.147 | -0.244 | 0.161 | -0.270 | 0.129 |
| Riparian quality | OOPs | -0.008 | 0.020 | -0.038 | 0.684 | -0.005 | 0.002 | -0.108 | 0.049 | -0.013 | 0.019 | -0.146 | 0.511 |
| OOPs | Riparian width | -0.004 | 0.032 | -0.019 | 0.898 | -0.042 | 0.043 | -0.159 | 0.332 | -0.046 | 0.052 | -0.178 | 0.374 |
| VOPs | Riparian width | 0.042 | 0.063 | 0.123 | 0.511 | 0.047 | 0.045 | 0.117 | 0.292 | 0.089 | 0.043 | 0.240 | 0.038 |
| Riparian quality | Riparian width | 0.024 | 0.005 | 0.529 | 0.000 | 0.003 | 0.004 | 0.049 | 0.473 | 0.027 | 0.005 | 0.578 | 0.000 |
| OOPs | Age of palms | -0.275 | 0.154 | -0.471 | 0.073 | -0.001 | 0.005 | -0.006 | 0.899 | -0.276 | 0.153 | -0.477 | 0.071 |
| VOPs | Age of palms | 0.259 | 0.143 | 0.288 | 0.071 | 0.054 | 0.058 | 0.094 | 0.358 | 0.313 | 0.120 | 0.382 | 0.009 |

**Table S6** Direct and indirect effects between observed variables in the proposed causal model at **5 km scale forest cover *G. boninense* attack**. For more details, see Figure A3. Statistically significant (alpha level of 0.05) effects/coefficients are in bold letters. b = Unstandardized effects. b* = Standardised effects/ coefficients. SE = Standard error. p = p-value.

|  |  | **Direct effect** | | | | **Indirect effect** | | | | **Total effect** | | | |
| --- | --- | --- | --- | --- | --- | --- | --- | --- | --- | --- | --- | --- | --- |
| **Affected variable** | **Causal variable** | ***b*** | ***SE*** | ***b**** | ***p*** | ***b*** | ***SE*** | ***b**** | ***p*** | ***b*** | ***SE*** | ***b**** | ***p*** |
| *Ganoderma* fungus attack | Riparian width | -0.001 | 0.000 | -0.044 | 0.014 | 0.000 | 0.000 | 0.015 | 0.200 | -0.001 | 0.000 | -0.029 | 0.152 |
| *Ganoderma* fungus attack | Riparian quality | 0.009 | 0.011 | 0.016 | 0.412 | 0.000 | 0.001 | -0.014 | 0.615 | 0.010 | 0.011 | 0.002 | 0.391 |
| *Ganoderma* fungus attack | VOPs | -0.001 | 0.001 | -0.011 | 0.522 | 0.001 | 0.001 | 0.012 | 0.085 | 0.000 | 0.001 | 0.001 | 0.807 |
| *Ganoderma* fungus attack | OOPs | -0.003 | 0.002 | -0.028 | 0.125 | 0.000 | 0.001 | 0.026 | 0.910 | -0.003 | 0.002 | -0.002 | 0.080 |
| *Ganoderma* fungus attack | Distance to nearest forest | -0.001 | 0.001 | -0.011 | 0.259 | -0.004 | 0.003 | -0.005 | 0.199 | -0.005 | 0.002 | -0.016 | 0.037 |
| *Ganoderma* fungus attack | Distance from riparian edge | 0.001 | 0.001 | 0.005 | 0.205 | 0.000 | - | 0.000 | - | 0.001 | 0.001 | 0.005 | 0.205 |
| *Ganoderma* fungus attack | Forest cover | 0.006 | 0.003 | 0.040 | 0.080 | 0.000 | 0.001 | -0.031 | 0.679 | 0.006 | 0.004 | 0.009 | 0.070 |
| *Ganoderma* fungus attack | Age of palms | 0.003 | 0.001 | 0.041 | 0.005 | -0.001 | 0.001 | -0.021 | 0.082 | 0.002 | 0.001 | 0.020 | 0.002 |
| Riparian quality | Forest cover | 0.004 | 0.062 | 0.017 | 0.945 | 0.034 | 0.024 | -0.160 | 0.150 | 0.038 | 0.060 | -0.143 | 0.525 |
| Riparian quality | Distance to nearest forest | -0.065 | 0.029 | -0.295 | 0.028 | -0.001 | 0.033 | 0.137 | 0.972 | -0.066 | 0.045 | -0.158 | 0.142 |
| VOPs | OOPs | -0.171 | 0.170 | -0.111 | 0.314 | -0.076 | 0.050 | -0.149 | 0.124 | -0.247 | 0.162 | -0.260 | 0.127 |
| Riparian quality | OOPs | 0.003 | 0.038 | 0.013 | 0.946 | 0.001 | 0.006 | -0.126 | 0.863 | 0.004 | 0.033 | -0.113 | 0.915 |
| OOPs | Riparian width | 0.016 | 0.038 | 0.07 | 0.680 | -0.045 | 0.048 | -0.170 | 0.351 | -0.029 | 0.056 | -0.100 | 0.603 |
| VOPs | Riparian width | 0.041 | 0.064 | 0.123 | 0.512 | 0.045 | 0.045 | 0.108 | 0.327 | 0.086 | 0.043 | 0.231 | 0.047 |
| Riparian quality | Riparian width | 0.028 | 0.005 | 0.624 | 0.000 | -0.002 | 0.002 | -0.028 | 0.471 | 0.026 | 0.005 | 0.596 | 0.000 |
| OOPs | Age of palms | -0.292 | 0.153 | -0.501 | 0.055 | 0.002 | 0.006 | 0.024 | 0.708 | -0.290 | 0.152 | -0.477 | 0.056 |
| VOPs | Age of palms | 0.259 | 0.146 | 0.288 | 0.076 | 0.056 | 0.064 | 0.094 | 0.379 | 0.315 | 0.119 | 0.382 | 0.008 |

**Table S7** Direct and indirect effects between observed variables in the proposed causal model at **10 km scale forest cover *G. boninense* attack**. For more details, see Figure A3. Statistically significant (alpha level of 0.05) effects/coefficients are in bold letters. b = Unstandardized effects. b* = Standardised effects/ coefficients. SE = Standard error. p = p-value.

|  |  | **Direct effect** | | | | **Indirect effect** | | | | **Total effect** | | | |
| --- | --- | --- | --- | --- | --- | --- | --- | --- | --- | --- | --- | --- | --- |
| **Affected variable** | **Causal variable** | ***b*** | ***SE*** | ***b**** | ***p*** | ***b*** | ***SE*** | ***b**** | ***p*** | ***b*** | ***SE*** | ***b**** | ***p*** |
| *Ganoderma* fungus attack | Riparian width | -0.001 | 0.000 | -0.040 | 0.017 | 0.000 | 0.000 | 0.014 | 0.216 | -0.001 | 0.000 | -0.025 | 0.187 |
| *Ganoderma* fungus attack | Riparian quality | 0.010 | 0.012 | 0.017 | 0.382 | 0.001 | 0.001 | -0.013 | 0.655 | 0.011 | 0.011 | 0.004 | 0.341 |
| *Ganoderma* fungus attack | VOPs | -0.001 | 0.001 | -0.012 | 0.471 | 0.001 | 0.001 | 0.014 | 0.083 | 0.000 | 0.001 | 0.002 | 0.881 |
| *Ganoderma* fungus attack | OOPs | -0.003 | 0.002 | -0.026 | 0.138 | 0.000 | 0.001 | 0.022 | 0.888 | -0.003 | 0.002 | -0.004 | 0.089 |
| *Ganoderma* fungus attack | Distance to nearest forest | -0.001 | 0.001 | -0.007 | 0.487 | -0.011 | 0.008 | -0.009 | 0.162 | -0.012 | 0.007 | -0.017 | 0.099 |
| *Ganoderma* fungus attack | Distance from riparian edge | 0.001 | 0.001 | 0.005 | 0.205 | 0.000 | - | 0.000 | - | 0.001 | 0.001 | 0.005 | 0.205 |
| *Ganoderma* fungus attack | Forest cover | 0.011 | 0.006 | 0.042 | 0.088 | 0.001 | 0.001 | -0.027 | 0.415 | 0.012 | 0.007 | 0.014 | 0.066 |
| *Ganoderma* fungus attack | Age of palms | 0.003 | 0.001 | 0.038 | 0.008 | -0.001 | 0.001 | -0.017 | 0.116 | 0.002 | 0.001 | 0.021 | 0.003 |
| Riparian quality | Forest cover | -0.005 | 0.109 | -0.013 | 0.959 | 0.065 | 0.041 | -0.103 | 0.113 | 0.060 | 0.106 | -0.116 | 0.572 |
| Riparian quality | Distance to nearest forest | -0.067 | 0.028 | -0.304 | 0.019 | 0.007 | 0.108 | 0.133 | 0.951 | -0.060 | 0.109 | -0.171 | 0.585 |
| VOPs | OOPs | -0.171 | 0.171 | -0.112 | 0.318 | -0.079 | 0.050 | -0.142 | 0.118 | -0.250 | 0.163 | -0.254 | 0.125 |
| Riparian quality | OOPs | 0.005 | 0.037 | 0.027 | 0.883 | 0.001 | 0.005 | -0.124 | 0.913 | 0.006 | 0.032 | -0.097 | 0.852 |
| OOPs | Riparian width | 0.030 | 0.039 | 0.130 | 0.459 | -0.047 | 0.051 | -0.175 | 0.358 | -0.017 | 0.061 | -0.045 | 0.773 |
| VOPs | Riparian width | 0.041 | 0.064 | 0.122 | 0.517 | 0.043 | 0.047 | 0.103 | 0.364 | 0.084 | 0.044 | 0.225 | 0.053 |
| Riparian quality | Riparian width | 0.027 | 0.005 | 0.616 | 0.000 | -0.001 | 0.002 | -0.022 | 0.568 | 0.026 | 0.005 | 0.594 | 0.000 |
| OOPs | Age of palms | -0.304 | 0.154 | -0.517 | 0.048 | 0.004 | 0.007 | 0.044 | 0.547 | -0.300 | 0.153 | -0.473 | 0.050 |
| VOPs | Age of palms | 0.259 | 0.148 | 0.287 | 0.080 | 0.058 | 0.067 | 0.095 | 0.389 | 0.317 | 0.118 | 0.382 | 0.007 |

**Table S8** Direct and indirect effects between observed variables in the proposed causal model at **20 km scale forest cover *G. boninense* attack**. For more details, see Figure A3. Statistically significant (alpha level of 0.05) effects/coefficients are in bold letters. b = Unstandardized effects. b* = Standardised effects/ coefficients. SE = Standard error. p = p-value.

|  |  | **Direct effect** | | | | **Indirect effect** | | | | **Total effect** | | | |
| --- | --- | --- | --- | --- | --- | --- | --- | --- | --- | --- | --- | --- | --- |
| **Affected variable** | **Causal variable** | ***b*** | ***SE*** | ***b**** | ***p*** | ***b*** | ***SE*** | ***b**** | ***p*** | ***b*** | ***SE*** | ***b**** | ***p*** |
| *Ganoderma* fungus attack | Riparian width | -0.002 | 0.000 | -0.059 | 0.001 | 0.001 | 0.000 | 0.030 | 0.069 | -0.001 | 0.001 | -0.029 | 0.177 |
| *Ganoderma* fungus attack | Riparian quality | 0.021 | 0.014 | 0.037 | 0.125 | -0.000 | 0.001 | -0.034 | 0.423 | 0.021 | 0.013 | 0.003 | 0.123 |
| *Ganoderma* fungus attack | VOPs | -0.002 | 0.001 | -0.024 | 0.147 | 0.001 | 0.000 | 0.034 | 0.006 | -0.001 | 0.001 | 0.009 | 0.546 |
| *Ganoderma* fungus attack | OOPs | -0.000 | 0.001 | -0.004 | 0.706 | -0.000 | 0.001 | -0.020 | 0.574 | -0.001 | 0.001 | -0.024 | 0.410 |
| *Ganoderma* fungus attack | Distance to nearest forest | -0.004 | 0.001 | -0.029 | 0.000 | 0.002 | 0.002 | -0.001 | 0.409 | -0.002 | 0.002 | -0.030 | 0.442 |
| *Ganoderma* fungus attack | Distance from riparian edge | 0.001 | 0.001 | 0.005 | 0.205 | 0.000 | - | 0.000 | - | 0.001 | 0.001 | 0.005 | 0.205 |
| *Ganoderma* fungus attack | Forest cover | 0.008 | 0.003 | 0.040 | 0.013 | -0.004 | 0.002 | -0.030 | 0.100 | 0.004 | 0.003 | 0.010 | 0.159 |
| *Ganoderma* fungus attack | Age of palms | 0.003 | 0.001 | 0.046 | 0.000 | -0.001 | 0.001 | -0.014 | 0.193 | 0.002 | 0.001 | 0.032 | 0.001 |
| Riparian quality | Forest cover | -0.066 | 0.055 | -0.193 | 0.235 | -0.026 | 0.026 | -0.219 | 0.312 | -0.092 | 0.054 | -0.412 | 0.090 |
| Riparian quality | Distance to nearest forest | -0.051 | 0.030 | -0.223 | 0.086 | -0.033 | 0.039 | -0.113 | 0.400 | -0.084 | 0.040 | -0.336 | 0.037 |
| VOPs | OOPs | -0.171 | 0.170 | -0.111 | 0.313 | -0.076 | 0.050 | -0.150 | 0.128 | -0.247 | 0.161 | -0.261 | 0.126 |
| Riparian quality | OOPs | -0.002 | 0.020 | -0.006 | 0.949 | -0.004 | 0.003 | -0.195 | 0.113 | -0.006 | 0.020 | -0.201 | 0.778 |
| OOPs | Riparian width | 0.013 | 0.038 | 0.061 | 0.727 | -0.044 | 0.048 | -0.169 | 0.355 | -0.031 | 0.053 | -0.108 | 0.557 |
| VOPs | Riparian width | 0.042 | 0.063 | 0.123 | 0.511 | 0.045 | 0.044 | 0.109 | 0.311 | 0.087 | 0.044 | 0.232 | 0.049 |
| Riparian quality | Riparian width | 0.027 | 0.005 | 0.573 | 0.000 | 0.001 | 0.002 | 0.026 | 0.475 | 0.028 | 0.005 | 0.599 | 0.000 |
| OOPs | Age of palms | -0.291 | 0.151 | -0.497 | 0.054 | 0.002 | 0.007 | 0.020 | 0.755 | -0.289 | 0.150 | -0.477 | 0.055 |
| VOPs | Age of palms | 0.259 | 0.146 | 0.288 | 0.076 | 0.056 | 0.062 | 0.094 | 0.372 | 0.315 | 0.119 | 0.382 | 0.008 |

**Table S9** Direct and indirect effects between observed variables in the proposed causal model at **30 km scale forest cover *G. boninense* attack**. For more details, see Figure A3. Statistically significant (alpha level of 0.05) effects/coefficients are in bold letters. b = Unstandardized effects. b* = Standardised effects/ coefficients. SE = Standard error. p = p-value.

|  |  | **Direct effect** | | | | **Indirect effect** | | | | **Total effect** | | | |
| --- | --- | --- | --- | --- | --- | --- | --- | --- | --- | --- | --- | --- | --- |
| **Affected variable** | **Causal variable** | ***b*** | ***SE*** | ***b**** | ***p*** | ***b*** | ***SE*** | ***b**** | ***p*** | ***b*** | ***SE*** | ***b**** | ***p*** |
| *Ganoderma* fungus attack | Riparian width | -0.001 | 0.001 | -0.050 | 0.024 | 0.001 | 0.001 | 0.027 | 0.161 | -0.001 | 0.001 | -0.024 | 0.400 |
| *Ganoderma* fungus attack | Riparian quality | 0.015 | 0.013 | 0.027 | 0.243 | -0.000 | 0.000 | -0.019 | 0.581 | 0.015 | 0.013 | 0.008 | 0.241 |
| *Ganoderma* fungus attack | VOPs | -0.002 | 0.002 | -0.019 | 0.358 | 0.001 | 0.000 | 0.029 | 0.013 | -0.000 | 0.001 | 0.010 | 0.838 |
| *Ganoderma* fungus attack | OOPs | 0.000 | 0.001 | -0.002 | 0.855 | -0.001 | 0.001 | -0.014 | 0.301 | -0.001 | 0.001 | -0.016 | 0.450 |
| *Ganoderma* fungus attack | Distance to nearest forest | -0.005 | 0.001 | -0.040 | 0.000 | 0.005 | 0.002 | 0.006 | 0.016 | 0.000 | 0.002 | -0.034 | 0.993 |
| *Ganoderma* fungus attack | Distance from riparian edge | 0.001 | 0.001 | 0.005 | 0.205 | 0.000 | - | 0.000 | - | 0.001 | 0.001 | 0.005 | 0.205 |
| *Ganoderma* fungus attack | Forest cover | 0.007 | 0.002 | 0.032 | 0.001 | -0.008 | 0.003 | -0.049 | 0.004 | -0.001 | 0.002 | -0.016 | 0.542 |
| *Ganoderma* fungus attack | Age of palms | 0.003 | 0.001 | 0.048 | 0.000 | -0.000 | 0.000 | -0.010 | 0.389 | 0.003 | 0.001 | 0.037 | 0.000 |
| Riparian quality | Forest cover | -0.108 | 0.059 | -0.289 | 0.069 | -0.028 | 0.046 | -0.207 | 0.542 | -0.136 | 0.044 | -0.497 | 0.002 |
| Riparian quality | Distance to nearest forest | -0.024 | 0.040 | -0.105 | 0.548 | -0.122 | 0.078 | -0.290 | 0.116 | -0.146 | 0.063 | -0.395 | 0.020 |
| VOPs | OOPs | -0.171 | 0.169 | -0.111 | 0.312 | -0.073 | 0.051 | -0.159 | 0.147 | -0.244 | 0.161 | -0.270 | 0.129 |
| Riparian quality | OOPs | -0.008 | 0.020 | -0.038 | 0.684 | -0.005 | 0.002 | -0.108 | 0.049 | -0.013 | 0.019 | -0.146 | 0.511 |
| OOPs | Riparian width | -0.004 | 0.032 | -0.019 | 0.898 | -0.042 | 0.043 | -0.159 | 0.332 | -0.046 | 0.052 | -0.178 | 0.374 |
| VOPs | Riparian width | 0.042 | 0.063 | 0.123 | 0.511 | 0.047 | 0.045 | 0.117 | 0.292 | 0.089 | 0.043 | 0.240 | 0.038 |
| Riparian quality | Riparian width | 0.024 | 0.005 | 0.529 | 0.000 | 0.003 | 0.004 | 0.049 | 0.473 | 0.027 | 0.005 | 0.578 | 0.000 |
| OOPs | Age of palms | -0.275 | 0.154 | -0.471 | 0.073 | -0.001 | 0.005 | -0.006 | 0.899 | -0.276 | 0.153 | -0.477 | 0.071 |
| VOPs | Age of palms | 0.259 | 0.143 | 0.288 | 0.071 | 0.054 | 0.058 | 0.094 | 0.358 | 0.313 | 0.120 | 0.382 | 0.009 |

**Table S10** Direct and indirect effects between observed variables in the proposed causal model at **5 km scale forest cover lepidopteran caterpillar attack**. For more details, see Figure A3. Statistically significant (alpha level of 0.05) effects/coefficients are in bold letters. b = Unstandardized effects. b* = Standardised effects / coefficients. SE = Standard error. p = p-value.

|  |  | **Direct effect** | | | | **Indirect effect** | | | | **Total effect** | | | |
| --- | --- | --- | --- | --- | --- | --- | --- | --- | --- | --- | --- | --- | --- |
| **Affected variable** | **Causal variable** | ***b*** | ***SE*** | ***b**** | ***p*** | ***b*** | ***SE*** | ***b**** | ***p*** | ***b*** | ***SE*** | ***b**** | ***p*** |
| Caterpillar attack | Riparian width | 0.035 | **0.021** | 0.076 | 0.095 | -0.042 | 0.074 | -0.071 | 0.570 | -0.007 | 0.076 | 0.006 | 0.924 |
| Caterpillar attack | Riparian quality | 0.307 | 1.073 | 0.030 | 0.775 | 0.002 | 0.014 | -0.025 | 0.865 | 0.310 | 1.063 | 0.005 | 0.771 |
| Caterpillar attack | VOPs | 0.236 | 0.074 | 0.176 | 0.002 | -0.156 | 0.052 | -0.113 | 0.003 | 0.080 | 0.100 | 0.064 | 0.426 |
| Caterpillar attack | OOPs | 0.084 | 0.125 | 0.041 | 0.499 | 0.072 | 0.074 | 0.128 | 0.332 | 0.156 | 0.133 | 0.169 | 0.240 |
| Caterpillar attack | Distance to nearest forest | 0.039 | 0.146 | 0.018 | 0.788 | -0.102 | 0.132 | -0.092 | 0.437 | -0.063 | 0.148 | -0.074 | 0.669 |
| Caterpillar attack | Distance from riparian edge | -0.079 | 0.051 | -0.023 | 0.123 | 0.000 | 0.000 | 0.000 | 0.002 | -0.079 | 0.051 | -0.023 | 0.123 |
| Caterpillar attack | Forest cover | 0.056 | 0.212 | 0.022 | 0.790 | 0.080 | 0.059 | 0.147 | 0.176 | 0.136 | 0.219 | 0.169 | 0.533 |
| Caterpillar attack | Age of palms | -0.437 | 0.104 | -0.363 | 0.000 | 0.047 | 0.042 | 0.083 | 0.266 | -0.390 | 0.090 | -0.280 | 0.000 |
| Riparian quality | Forest cover | 0.004 | 0.062 | 0.017 | 0.945 | 0.034 | 0.024 | -0.160 | 0.150 | 0.038 | 0.060 | -0.143 | 0.525 |
| Riparian quality | Distance to nearest forest | -0.064 | 0.029 | -0.295 | 0.028 | -0.001 | 0.033 | 0.137 | 0.972 | -0.066 | 0.045 | -0.158 | 0.142 |
| VOPs | OOPs | -0.171 | 0.170 | -0.111 | 0.314 | -0.076 | 0.050 | -0.149 | 0.124 | -0.247 | 0.162 | -0.260 | 0.127 |
| Riparian quality | OOPs | 0.003 | 0.038 | 0.013 | 0.946 | 0.001 | 0.006 | -0.126 | 0.863 | 0.004 | 0.033 | -0.113 | 0.915 |
| OOPs | Riparian width | 0.015 | 0.038 | 0.070 | 0.680 | -0.045 | 0.048 | -0.170 | 0.351 | -0.029 | 0.056 | -0.100 | 0.603 |
| VOPs | Riparian width | 0.042 | 0.064 | 0.123 | 0.512 | 0.045 | 0.045 | 0.108 | 0.327 | 0.086 | 0.043 | 0.231 | 0.047 |
| Riparian quality | Riparian width | 0.028 | 0.005 | 0.624 | 0.000 | -0.002 | 0.002 | -0.028 | 0.471 | 0.026 | 0.005 | 0.596 | 0.000 |
| OOPs | Age of palms | -0.293 | 0.153 | -0.501 | 0.055 | 0.002 | 0.006 | 0.024 | 0.708 | -0.290 | 0.152 | -0.477 | 0.056 |
| VOPs | Age of palms | 0.259 | 0.146 | 0.288 | 0.076 | 0.056 | 0.064 | 0.094 | 0.379 | 0.315 | 0.118 | 0.382 | 0.008 |

**Table S11** Direct and indirect effects between observed variables in the proposed causal model at **10 km scale forest cover lepidopteran caterpillar attack**. For more details, see Figure A3. Statistically significant (alpha level of 0.05) effects/coefficients are in bold letters. b = Unstandardized effects. b* = Standardised effects / coefficients. SE = Standard error. p = p-value.

|  |  | **Direct effect** | | | | **Indirect effect** | | | | **Total effect** | | | |
| --- | --- | --- | --- | --- | --- | --- | --- | --- | --- | --- | --- | --- | --- |
| **Affected variable** | **Causal variable** | ***b*** | ***SE*** | ***b**** | ***p*** | ***b*** | ***SE*** | ***b**** | ***p*** | ***b*** | ***SE*** | ***b**** | ***p*** |
| Caterpillar attack | Riparian width | 0.034 | 0.020 | 0.075 | 0.090 | -0.041 | 0.074 | -0.068 | 0.581 | -0.007 | 0.077 | 0.006 | 0.927 |
| Caterpillar attack | Riparian quality | 0.312 | 1.074 | 0.031 | 0.771 | 0.002 | 0.020 | -0.027 | 0.931 | 0.314 | 1.066 | 0.004 | 0.769 |
| Caterpillar attack | VOPs | 0.235 | 0.075 | 0.176 | 0.002 | -0.160 | 0.052 | -0.113 | 0.002 | 0.075 | 0.099 | 0.064 | 0.444 |
| Caterpillar attack | OOPs | 0.104 | 0.129 | 0.051 | 0.419 | 0.074 | 0.076 | 0.120 | 0.328 | 0.178 | 0.126 | 0.171 | 0.156 |
| Caterpillar attack | Distance to nearest forest | 0.029 | 0.148 | 0.013 | 0.845 | -0.083 | 0.360 | -0.083 | 0.818 | -0.054 | 0.354 | -0.070 | 0.879 |
| Caterpillar attack | Distance from riparian edge | -0.079 | 0.051 | -0.023 | 0.123 | -0.000 | 0.000 | -0.000 | 0.002 | -0.079 | 0.051 | -0.023 | 0.123 |
| Caterpillar attack | Forest cover | 0.016 | 0.339 | 0.003 | 0.963 | 0.038 | 0.099 | 0.143 | 0.700 | 0.054 | 0.358 | 0.147 | 0.880 |
| Caterpillar attack | Age of palms | -0.442 | 0.103 | -0.367 | 0.000 | 0.051 | 0.040 | 0.086 | 0.208 | -0.391 | 0.090 | -0.280 | 0.000 |
| Riparian quality | Forest cover | -0.006 | 0.109 | -0.013 | 0.459 | 0.065 | 0.041 | -0.103 | 0.113 | 0.060 | 0.106 | -0.116 | 0.572 |
| Riparian quality | Distance to nearest forest | -0.066 | 0.028 | -0.304 | 0.959 | 0.007 | 0.108 | 0.133 | 0.951 | -0.060 | 0.109 | -0.171 | 0.585 |
| VOPs | OOPs | -0.171 | 0.171 | -0.112 | 0.318 | -0.079 | 0.050 | -0.142 | 0.118 | -0.250 | 0.163 | -0.254 | 0.125 |
| Riparian quality | OOPs | 0.005 | 0.037 | 0.027 | 0.883 | 0.001 | 0.005 | -0.124 | 0.913 | 0.006 | 0.032 | -0.097 | 0.852 |
| OOPs | Riparian width | 0.029 | 0.039 | 0.130 | 0.459 | -0.047 | 0.051 | -0.175 | 0.358 | -0.017 | 0.061 | -0.045 | 0.773 |
| VOPs | Riparian width | 0.042 | 0.064 | 0.122 | 0.517 | 0.043 | 0.047 | 0.103 | 0.364 | 0.084 | 0.044 | 0.225 | 0.053 |
| Riparian quality | Riparian width | 0.027 | 0.005 | 0.616 | 0.000 | -0.001 | 0.002 | -0.022 | 0.568 | 0.026 | 0.005 | 0.594 | 0.000 |
| OOPs | Age of palms | -0.305 | 0.154 | -0.517 | 0.048 | 0.004 | 0.007 | 0.044 | 0.547 | -0.300 | 0.153 | -0.473 | 0.050 |
| VOPs | Age of palms | 0.259 | 0.148 | 0.287 | 0.080 | 0.058 | 0.067 | 0.095 | 0.389 | 0.317 | 0.118 | 0.382 | 0.007 |

**Table S12** Direct and indirect effects between observed variables in the proposed causal model at **20 km scale forest cover lepidopteran caterpillar attack**. For more details, see Figure A3. Statistically significant (alpha level of 0.05) effects/coefficients are in bold letters. b = Unstandardized effects. b* = Standardised effects / coefficients. SE = Standard error. p = p-value.

|  |  | **Direct effect** | | | | **Indirect effect** | | | | **Total effect** | | | |
| --- | --- | --- | --- | --- | --- | --- | --- | --- | --- | --- | --- | --- | --- |
| **Affected variable** | **Causal variable** | ***b*** | ***SE*** | ***b**** | ***p*** | ***b*** | ***SE*** | ***b**** | ***p*** | ***b*** | ***SE*** | ***b**** | ***p*** |
| Caterpillar attack | Riparian width | 0.033 | 0.020 | 0.073 | 0.090 | -0.042 | 0.074 | -0.070 | 0.574 | -0.008 | 0.076 | 0.004 | 0.914 |
| Caterpillar attack | Riparian quality | 0.318 | 1.145 | 0.033 | 0.781 | -0.007 | 0.021 | -0.041 | 0.730 | 0.310 | 1.129 | -0.009 | 0.783 |
| Caterpillar attack | VOPs | 0.235 | 0.085 | 0.176 | 0.006 | -0.159 | 0.049 | -0.109 | 0.001 | 0.077 | 0.102 | 0.067 | 0.455 |
| Caterpillar attack | OOPs | 0.107 | 0.115 | 0.052 | 0.350 | 0.067 | 0.075 | 0.114 | 0.373 | 0.174 | 0.098 | 0.166 | 0.076 |
| Caterpillar attack | Distance to nearest forest | 0.026 | 0.140 | 0.011 | 0.855 | -0.005 | 0.104 | 0.015 | 0.960 | 0.020 | 0.094 | 0.026 | 0.829 |
| Caterpillar attack | Distance from riparian edge | -0.079 | 0.051 | -0.023 | 0.123 | 0.000 | 0.000 | 0.000 | 0.006 | -0.079 | 0.051 | -0.023 | 0.123 |
| Caterpillar attack | Forest cover | 0.004 | 0.184 | 0.001 | 0.982 | 0.023 | 0.071 | 0.066 | 0.748 | 0.027 | 0.158 | 0.067 | 0.864 |
| Caterpillar attack | Age of palms | -0.442 | 0.093 | -0.367 | 0.000 | 0.052 | 0.044 | 0.085 | 0.233 | -0.389 | 0.086 | -0.281 | 0.000 |
| Riparian quality | Forest cover | -0.066 | 0.055 | -0.193 | 0.235 | -0.026 | 0.026 | -0.219 | 0.312 | -0.092 | 0.054 | -0.412 | 0.090 |
| Riparian quality | Distance to nearest forest | -0.052 | 0.030 | -0.224 | 0.086 | -0.033 | 0.039 | -0.113 | 0.400 | -0.084 | 0.040 | -0.336 | 0.037 |
| VOPs | OOPs | -0.171 | 0.170 | -0.111 | 0.313 | -0.076 | 0.050 | -0.150 | 0.128 | -0.247 | 0.161 | -0.261 | 0.126 |
| Riparian quality | OOPs | -0.001 | 0.020 | -0.006 | 0.949 | -0.004 | 0.003 | -0.195 | 0.113 | -0.006 | 0.020 | -0.201 | 0.778 |
| OOPs | Riparian width | 0.013 | 0.038 | 0.061 | 0.727 | -0.044 | 0.048 | -0.169 | 0.355 | -0.031 | 0.053 | -0.108 | 0.557 |
| VOPs | Riparian width | 0.042 | 0.063 | 0.123 | 0.511 | 0.045 | 0.044 | 0.109 | 0.311 | 0.087 | 0.044 | 0.232 | 0.049 |
| Riparian quality | Riparian width | 0.027 | 0.005 | 0.573 | 0.000 | 0.001 | 0.002 | 0.026 | 0.475 | 0.028 | 0.005 | 0.599 | 0.000 |
| OOPs | Age of palms | -0.291 | 0.151 | -0.497 | 0.054 | 0.002 | 0.007 | 0.020 | 0.755 | -0.289 | 0.150 | -0.477 | 0.055 |
| VOPs | Age of palms | 0.259 | 0.146 | 0.288 | 0.076 | 0.056 | 0.062 | 0.094 | 0.372 | 0.315 | 0.119 | 0.382 | 0.008 |

**Table S13** Direct and indirect effects between observed variables in the proposed causal model at **30 km scale forest cover lepidopteran caterpillar attack**. For more details, see Figure A3. Statistically significant (alpha level of 0.05) effects/coefficients are in bold letters. b = Unstandardized effects. b* = Standardised effects / coefficients. SE = Standard error. p = p-value.

|  |  | **Direct effect** | | | | **Indirect effect** | | | | **Total effect** | | | |
| --- | --- | --- | --- | --- | --- | --- | --- | --- | --- | --- | --- | --- | --- |
| **Affected variable** | **Causal variable** | ***b*** | ***SE*** | ***b**** | ***p*** | ***b*** | ***SE*** | ***b**** | ***p*** | ***b*** | ***SE*** | ***b**** | ***p*** |
| Caterpillar attack | Riparian width | 0.032 | 0.021 | 0.071 | 0.121 | -0.039 | 0.073 | -0.065 | 0.591 | -0.007 | 0.075 | 0.006 | 0.926 |
| Caterpillar attack | Riparian quality | 0.467 | 1.130 | 0.048 | 0.679 | -0.028 | 0.039 | -0.044 | 0.469 | 0.439 | 1.105 | 0.004 | 0.691 |
| Caterpillar attack | VOPs | 0.217 | 0.084 | 0.163 | 0.010 | -0.154 | 0.048 | -0.093 | 0.001 | 0.063 | 0.099 | 0.070 | 0.525 |
| Caterpillar attack | OOPs | 0.126 | 0.101 | 0.061 | 0.212 | 0.056 | 0.070 | 0.088 | 0.423 | 0.182 | 0.095 | 0.149 | 0.056 |
| Caterpillar attack | Distance to nearest forest | -0.039 | **0.103** | -0.017 | 0.705 | 0.198 | 0.215 | 0.072 | 0.358 | 0.159 | 0.209 | 0.054 | 0.447 |
| Caterpillar attack | Distance from riparian edge | -0.079 | 0.051 | -0.023 | 0.123 | 0.000 | 0.000 | 0.000 | 0.010 | -0.079 | 0.051 | -0.023 | 0.123 |
| Caterpillar attack | Forest cover | 0.197 | 0.234 | 0.054 | 0.400 | -0.066 | 0.126 | 0.043 | 0.599 | 0.131 | 0.189 | 0.097 | 0.488 |
| Caterpillar attack | Age of palms | -0.411 | 0.077 | -0.343 | 0.000 | 0.032 | 0.046 | 0.070 | 0.486 | -0.379 | 0.079 | -0.273 | 0.000 |
| Riparian quality | Forest cover | -0.108 | 0.059 | -0.29 | 0.069 | -0.028 | 0.046 | -0.207 | 0.542 | -0.136 | 0.044 | -0.497 | 0.002 |
| Riparian quality | Distance to nearest forest | -0.024 | 0.040 | -0.105 | 0.548 | -0.122 | 0.078 | -0.290 | 0.116 | -0.146 | 0.063 | -0.395 | 0.020 |
| VOPs | OOPs | -0.171 | 0.169 | -0.111 | 0.312 | -0.073 | 0.051 | -0.159 | 0.147 | -0.244 | 0.161 | -0.270 | 0.129 |
| Riparian quality | OOPs | -0.008 | 0.020 | -0.038 | 0.684 | -0.005 | 0.002 | -0.108 | 0.049 | -0.013 | 0.019 | -0.146 | 0.511 |
| OOPs | Riparian width | -0.004 | 0.032 | -0.019 | 0.898 | -0.042 | 0.043 | -0.159 | 0.332 | -0.046 | 0.052 | -0.178 | 0.374 |
| VOPs | Riparian width | 0.042 | 0.063 | 0.123 | 0.511 | 0.047 | 0.045 | 0.117 | 00.292 | 0.089 | 0.043 | 0.240 | 0.038 |
| Riparian quality | Riparian width | 0.024 | 0.005 | 0.529 | 0.000 | 0.003 | 0.004 | 0.049 | 0.473 | 0.027 | 0.005 | 0.578 | 0.000 |
| OOPs | Age of palms | -0.275 | 0.154 | -0.471 | 0.073 | -0.001 | 0.005 | -0.006 | 0.899 | -0.276 | 0.153 | -0.477 | 0.071 |
| VOPs | Age of palms | 0.259 | 0.143 | 0.288 | 0.071 | 0.054 | 0.058 | 0.094 | 0.358 | 0.313 | 0.120 | 0.382 | 0.009 |

| 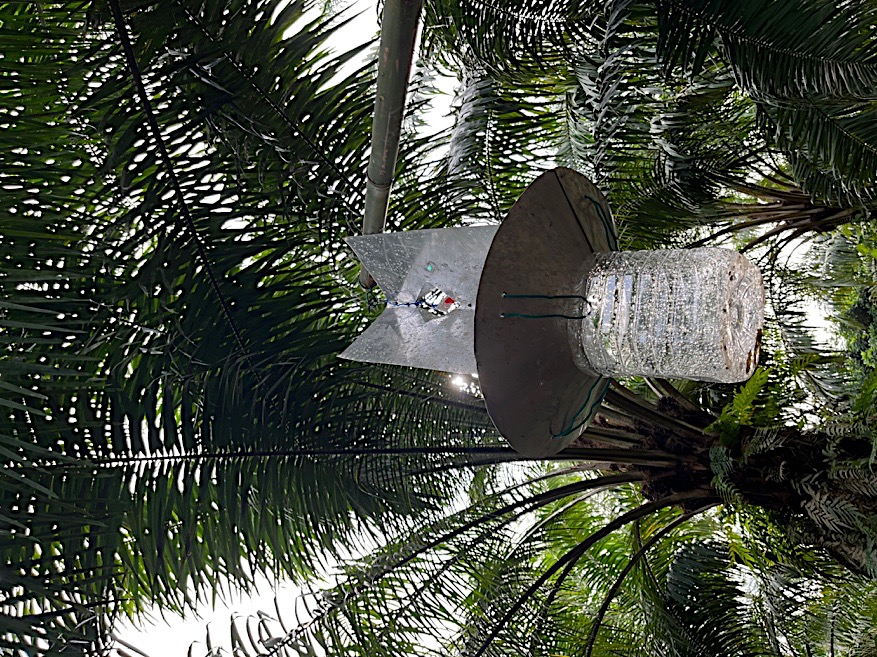  **Figure S1** Setup of pheromone trap in the field. | 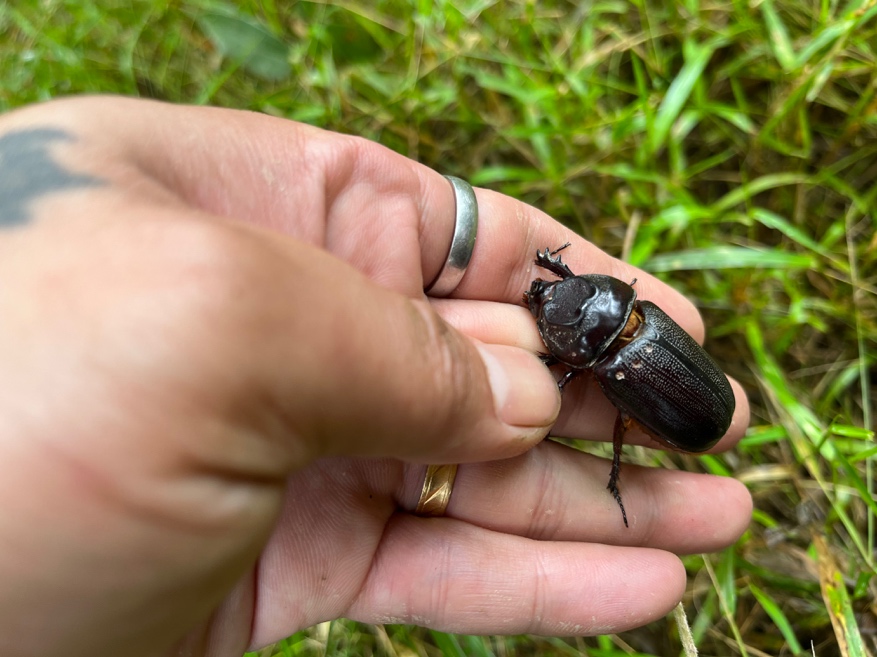  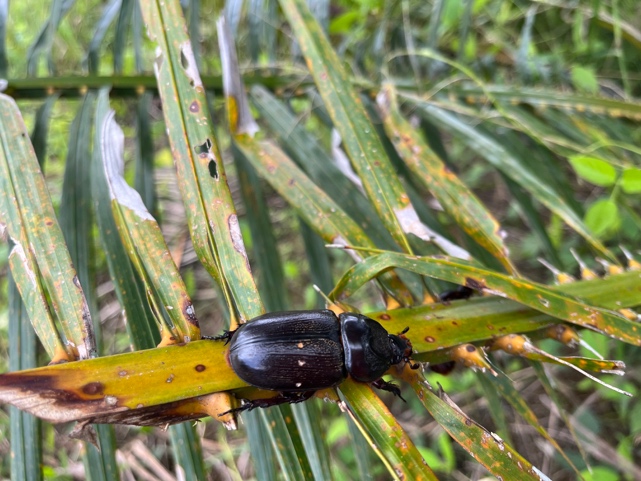  **Figure S2** Marked rhinoceros beetles with unique codes |
| --- | --- |
| 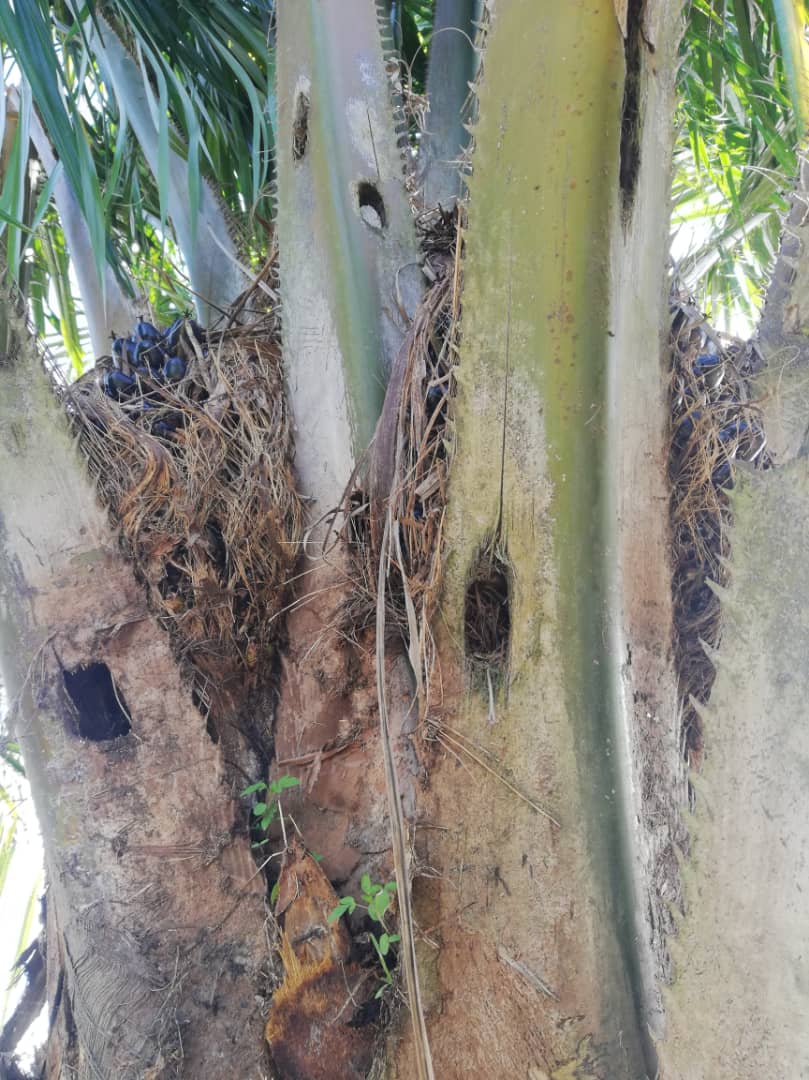  **Figure S3** Holes made by adult rhinoceros beetles in the base of fronds. | 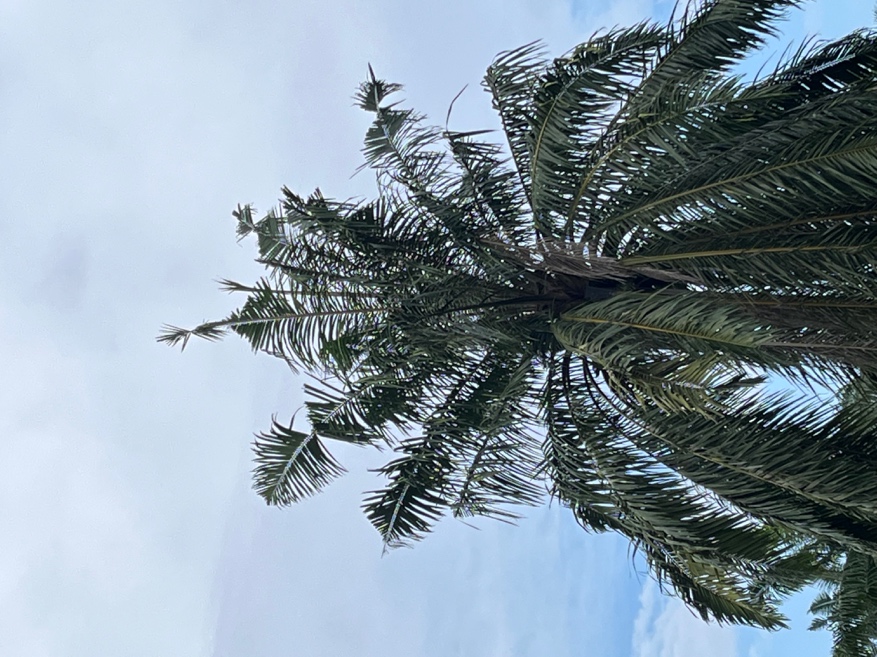  **Figure S4** The V-shaped cut damage caused by rhinoceros beetles as they tunnel into the crowns of mature palms. |
| 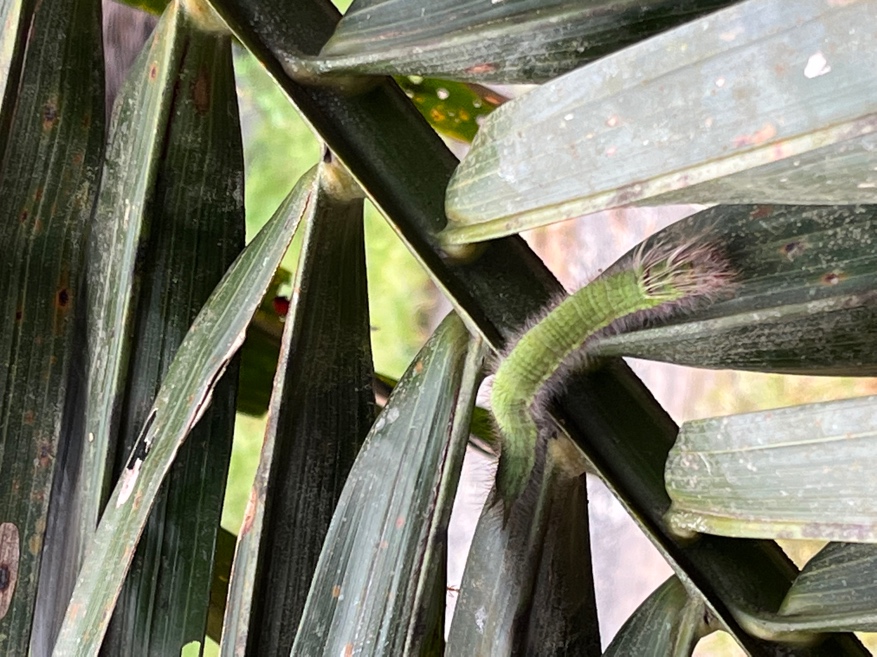  **Figure S5** Lepidopteran caterpillar on the palm leaves | 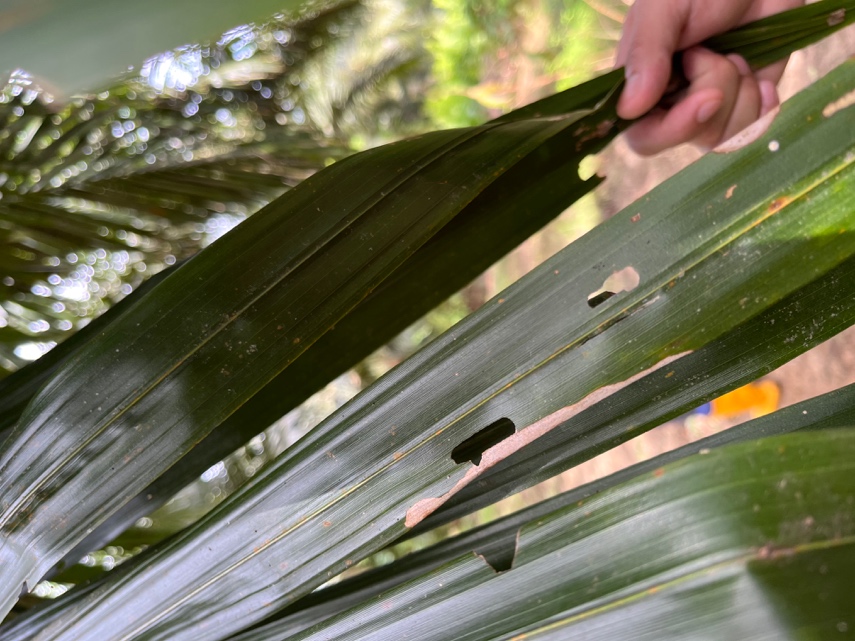  **Figure S6** Sign of attack from lepidopteran caterpillars |
| 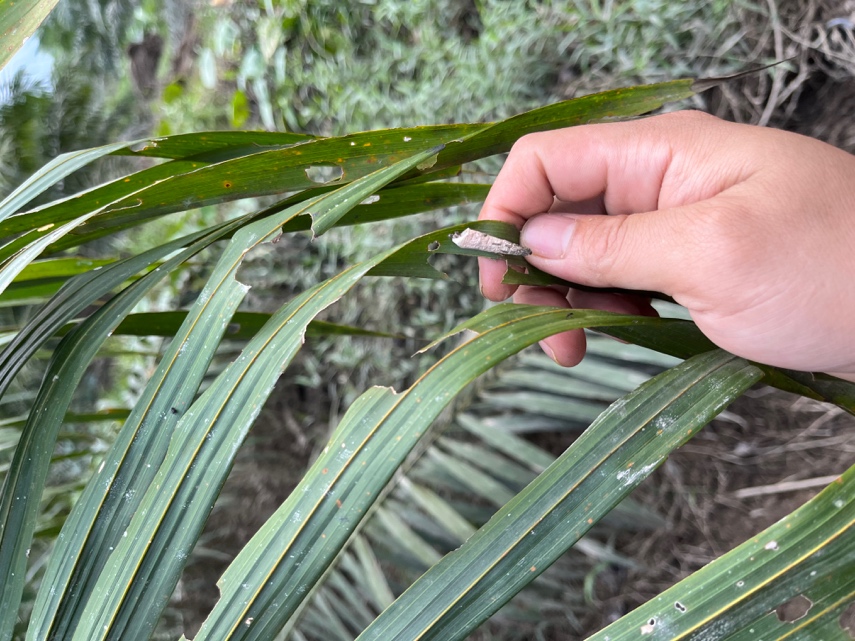  **Figure S7** Presence of bagworms on a palm leaf | 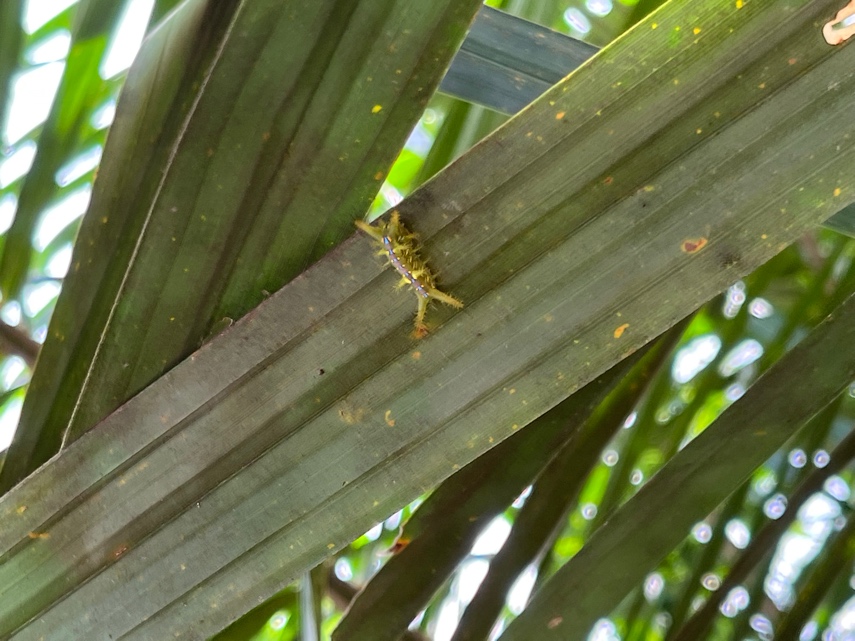  **Figure S8** Presence of nettle caterpillar on a palm leaf |
| 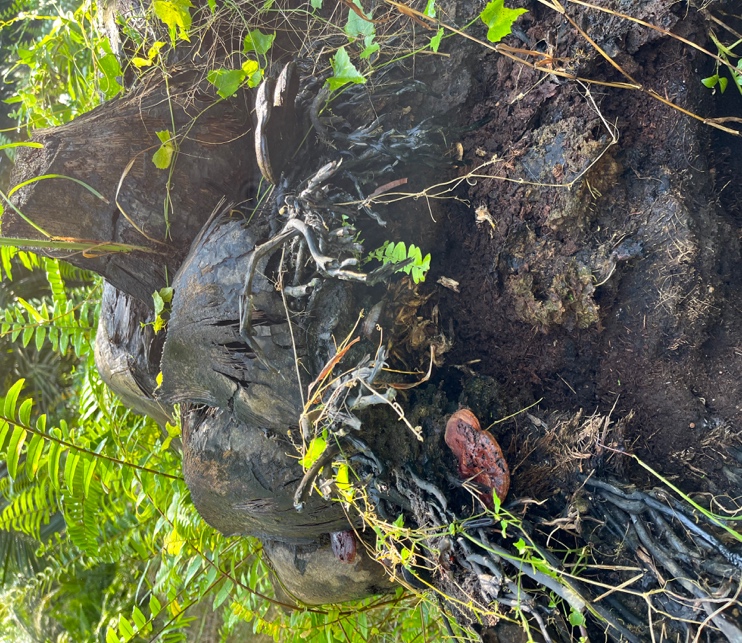  **Figure S9** Presence of fruiting body of *Ganoderma* fungus | 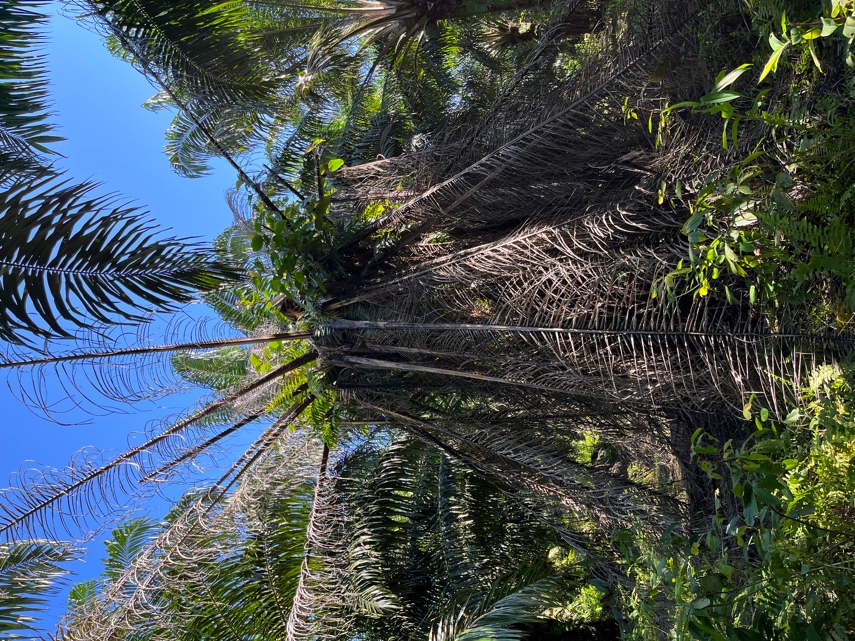  **Figure S10** Flattening of the crown (“skirting”) and unopened spear leaves caused by *Ganoderma* fungus |
